# Two Sides of the Same Coin: The Hippocampus as a Common Neural Substrate for Model-Based Planning and Spatial Memory

**DOI:** 10.1101/359232

**Authors:** Oliver Vikbladh, Michael R. Meager, John King, Karen Blackmon, Orrin Devinsky, Daphna Shohamy, Neil Burgess, Nathaniel D. Daw

**Affiliations:** Center for Neural Science, New York University, New York, NY, 10003 USA; Department of Psychology, New York University, New York, NY, 10003, USA; Department of Neurology, New York University School of Medicine, New York, NY, 10016, USA; Department of Clinical, Education & Health Psychology, Division of Psychology & Language Sciences, University College London, London, WC1H 0AP, UK; Department of Physiology, Neuroscience, and Behavioral Sciences, St. George’s University School of Medicine, St. George, Grenada, West Indies; Department of Neurosurgery, New York University School of Medicine, New York, NY, 10016, USA; Department of Psychiatry, New York University School of Medicine, New York, NY, 10016, USA; Department of Psychology and Zuckerman Mind, Brain, Behavior Institute, New York, NY, 10027, USA; Kavli Center for Brain Science, Columbia University, New York, NY, USA, 10027; Institute of Cognitive Neuroscience, University College London, London, WC1N 3AZ, UK; Institute of Neurology, University College London, London, WC1N 3BG, UK; Princeton Neuroscience Institute, Princeton University, Princeton, NJ, 08544, USA; Department of Psychology, Princeton University, Princeton, NJ, 08544, USA

## Abstract

Little is known about the neural mechanisms that allow humans and animals to plan actions using knowledge of task contingencies. Emerging theories hypothesize that it involves the same hippocampal mechanisms that support self-localization and memory for locations. Yet, there is limited direct evidence for the link between model-based planning and the hippocampal place map. We addressed this by investigating model-based planning and place memory in healthy controls and epilepsy patients treated using unilateral anterior temporal lobectomy with hippocampal resection. We found that both functions were impaired in the patient group. Specifically, the planning impairment was related to right hippocampal lesion size, controlling for overall lesion size. Furthermore, planning and place memory covaried with one another, but only in neurologically intact controls, consistent with both functions relying on the same structure in the healthy brain. These findings clarify the scope of hippocampal contributions to behavior and the neural mechanism of model-based planning.

## Introduction

Using knowledge of task contingencies, humans and other animals can plan novel courses of action, such as trajectories through a maze. Although the neural substrates for such “model-based” planning are poorly understood, this ability is often viewed as similar to other functions supported by the hippocampus, like representing and remembering locations in space. Both model-based planning (Tolman, 1948) and place memory (O’Keefe and Nadel, 1978) are often described as requiring ‘cognitive maps’ of the environment or task structure, and are contrasted against habitual response-based behaviors that depend on the basal ganglia. Still, despite the commonalities, these functions are distinct in principle and it is unclear whether they actually share a common neural mechanism and, if so, what that mechanism is.

Research into hippocampal spatial cognition most clearly emphasizes localization: determining one’s position in allocentric space. This function is most famously exemplified by location-selective neural responses in the hippocampus (O’Keefe and Nadel, 1978) and behaviorally operationalized using spatial tasks such as the Morris Water Maze (MWM) (Morris et al., 1982), where rodents have to find and remember the location of a hidden platform in an open arena. This type of allocentric “place memory” is distinguished from egocentric “response memory” (e.g., turn left or right) which relies on the basal ganglia (McDonald and White, 1994, Packard and McGaugh, 1996, Pearce et al., 1998). An analogous dissociation between “place-based” hippocampal and “response-based” basal ganglia-dependent memory, has been demonstrated in humans using functional magnetic resonance imaging (fMRI) in virtual spatial tasks (Hartley et al., 2003, Iaria et al., 2003, Voermans et al., 2004, Doeller et al., 2008).

By contrast, research into planning investigates how organisms use knowledge of task contingencies, like action outcomes and state transitions, to evaluate actions by mental simulation. Experiments probing such functions, including reward devaluation in operant lever pressing (Adams and Dickinson, 1981, Adams, 1982) and multi-step reinforcement learning (Gläscher et al., 2010, Daw et al., 2011), support a distinction between two classes of strategies – referred to as goal-directed or model-based planning vs. habitual or model-free learning (Balleine and Dickinson, 1998, Daw et al., 2005). This distinction seems to parallel the place vs. response memory dichotomy from spatial cognition (Poldrack and Packard, 2003, Kosaki et al., 2018) and the related declarative vs. procedural memory distinction from the memory literature (Squire, 1992, Knowlton et al., 1996, Foerde and Shohamy, 2011, Daw and Shohamy, 2015). Indeed, model-free learning, like egocentric response strategies in spatial tasks, is well captured by theories of dopamine and the basal ganglia (Schultz et al., 1997, Bayer and Glimcher, 2005).

It is less clear what neural mechanisms are responsible for model-based planning. There are, however, a number of suggestive reasons to suspect it shares a common hippocampal substrate with place memory (Hirsch, 1974, Dickinson and Balleine, 1993, Eilan et al., 1993, Johnson and Redish, 2007, Daw and Shohamy, 2015, Kumaran et al., 2016). Hippocampal function, of course, extends beyond spatial cognition to support declarative memory, and notably a role in encoding the relationships among environmental stimuli (Eichenbaum and Cohen, 2001, Davachi and Wagner, 2002, Kumaran et al., 2009, Schapiro et al., 2016, Boorman et al., 2016, Garvert et al., 2017). Knowing such relations is critical to building a model of task contingencies. Tests of relational encoding have even relied on tasks which are similar in logic to probes for model-based planning, like transitive inference or acquired equivalence (Dusek and Eichenbaum, 1997, Heckers et al., 2004, Shohamy and Wagner, 2008, Wimmer and Shohamy, 2012). Moreover, the hippocampus has been implicated in the ability to imagine or simulate future events, a function that may be critical to model-based planning (Hassibis et al., 2007, Addis et al., 2011). Spatial navigation studies have further demonstrated that the hippocampus and surrounding medial temporal areas, in addition to the current location, encode other variables that are relevant to planning, such as boundaries or the identity, direction, and distance to a goal (Spiers and Maguire, 2007, Viard et al., 2011, Chadwick et al., 2015, Wikenheiser and Redish, 2015, Brown et al., 2016, Kaplan et al., 2017). Non-local place-cell firing, such as preplay of locations ahead of the animal, has also been proposed to support planning by mental simulation of candidate routes, drawing on a cognitive model or map of the world (Johnson and Redish, 2007, Pfeiffer and Foster, 2013, Daw and Shohamy, 2015, Mattar and Daw, 2017).

At the same time, there is a surprising lack of direct evidence for hippocampal involvement in model-based planning. For predominant rodent models of model-based behavior, including outcome devaluation and contingency degradation in operant lever-pressing, hippocampal lesions have negligible effects (Corbit and Balleine, 2000, Corbit et al., 2002). One exception is a recent rodent study in which hippocampal lesions impaired model-based planning in a multi-step decision task (Miller et al., 2017). However, the extensive training needed to teach animals such sequential decision tasks may elicit model-free strategies that only mimic the signatures of planning in more lightly trained humans (Akam et al., 2015, Economides et al., 2015). Even seemingly model-based rodent behavior (Miller et al., 2017) could thus rely on the hippocampus for different reasons. Finally, with few exceptions (Simon and Daw, 2011), little evidence links hippocampal activity in human neuroimaging, or rodent place cell preplay to planning in tasks specifically designed to identify choice strategies that require knowledge of task contingencies.

We therefore sought to directly test the hypothesis that model-based planning shares a common hippocampal mechanism with place memory in humans. To this end, we studied the performance on a model-based planning task (Daw et al., 2011) and a spatial memory task (Doeller et al., 2008) in healthy controls and patients with medically intractable epilepsy, treated by unilateral anterior temporal lobectomy (ATL) with hippocampal resection. We investigated whether damage to the temporal lobe impaired model-based planning and place memory and how it affected the relationship between them. If the hippocampus is a common neural substrate for both functions, we expected hippocampal damage to impair performance on both tasks, that performance in each task should be correlated in individuals with functioning hippocampus, and that this correlation should itself be affected by hippocampal damage. Finally, because the lesions also affected overlying cortex, we explored to what extent performance in either task was related specifically to the extent of damage to hippocampus on either side, controlling for the overall extent of the lesion.

## Results

### Participant Characteristics

We recruited 19 epilepsy patients, treated with unilateral anterior temporal lobectomy (ATL) i.e. surgical removal of the anterior temporal lobe on one side (Table 1), and 19 healthy controls (see Methods). Patients and controls displayed no significant group differences in IQ (t_36_= 0.2200, p=0.8271), age (t_36_=−0.7760, p=0.4428) or number of males vs. females (z=−0.3261, p=0.7444).

**Table 1:**
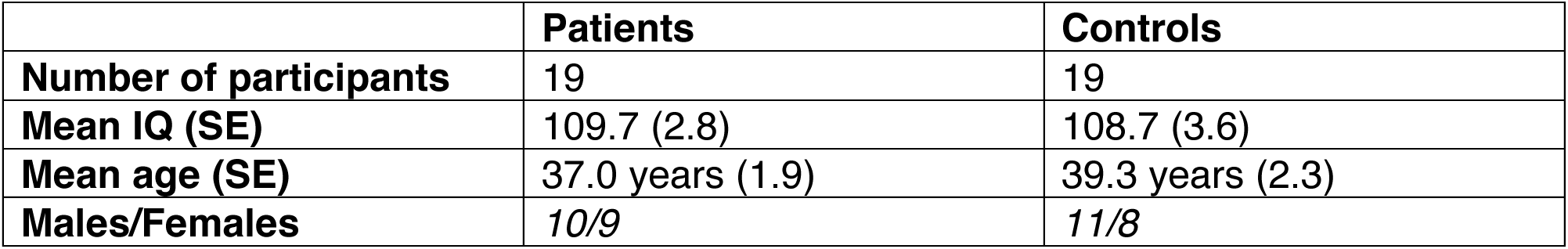
Characteristics of patients and controls.

For 10 out of 19 patients ATL was right lateralized (Table 2). There were no significant differences between the right and left lateralized ATL groups in IQ (t= −0.5295, p= 0.6033), age (t_17_=1.0876, p=0.2919) or number of males vs. females (z=−0.6752, p=0.4995). Participant-wise lesion masks were normalized to the MNI template (see Figure 1) and compared to the Harvard-Oxford Lexicon (p>.5) in order to estimate size of lesion to the hippocampus (See Methods). There was no group difference in estimated size of hippocampal resection (t_17_= − 1.1124, p= 0.2814), however, total lesion size was significantly larger in the right lateralized group (t_17_=−2.6103, p=0.0183).

**Table 2:**
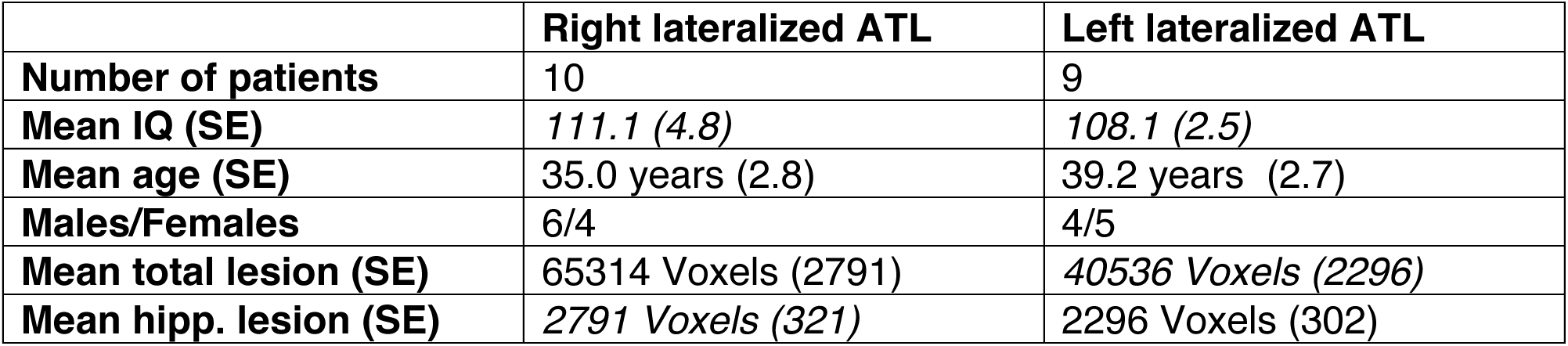
Characteristics for patients with right and left lateralized ATL.

|  | **Right lateralized ATL** | **Left lateralized ATL** |
| --- | --- | --- |
| **Number of patients** | 10 | 9 |
| **Mean IQ (SE)** | 111.1 (4.8) | 108.1 (2.5) |
| **Mean age (SE)** | 35.0 years (2.8) | 39.2 years (2.7) |
| **Males/Females** | 6/4 | 4/5 |
| **Mean total lesion (SE)** | 65314 Voxels (2791) | 40536 Voxels (2296) |
| **Mean hipp. lesion (SE)** | 2791 Voxels (321) | 2296 Voxels (302) |

**Figure 1:**
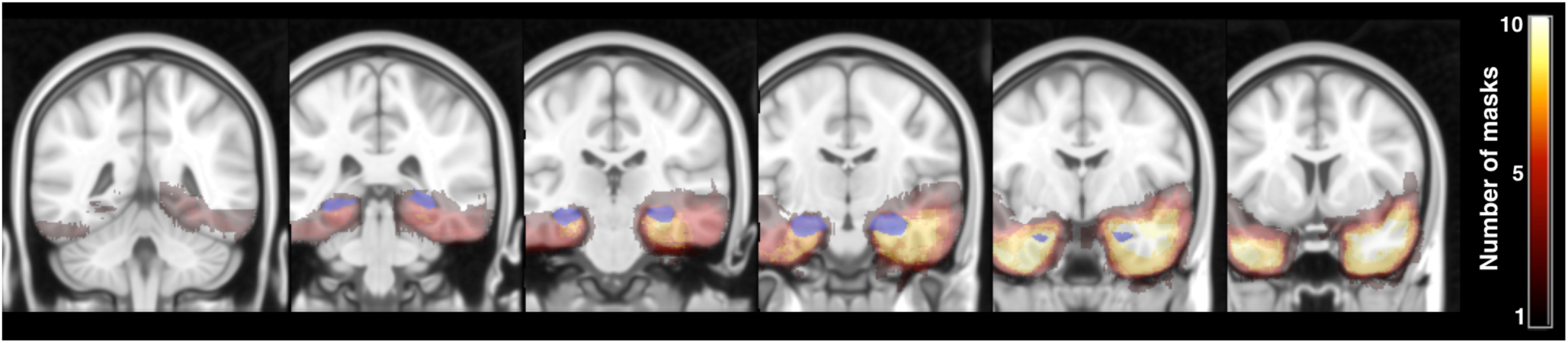
Patient lesion masks. Slices (y=82, 92, 102, 112, 122, 132) showing all 19 hand-drawn patient ATL lesion masks normalized to the MNI template. Heat maps indicate the number of masks overlapping at a given voxel. The hippocampus, as defined by Harvard-Oxford Lexicon (p>.5), is shown in blue.

### Patients display impaired place memory

Participants completed 64 trials of a spatial memory task (Doeller et al., 2008) where on each trial they had to indicate the correct location of one of four objects in a virtual arena (see Methods). For two objects (boundary objects), correct locations were defined with in relation to distal cues and the boundaries of the arena. For two objects (landmark objects) correct locations were defined in relation to a landmark in the arena (see Figure 2 Left). Participants were not instructed of the difference between landmark and boundary objects. Participants were also not aware that trials were presented in 4 blocks, between which the landmark moved in relation to the boundaries of the arena (see Figure 2 Left).

The mean number of completed trials was 61.3 (SE 0.5) with no significant difference between control and patient group in number of completed trials (t_36_−0.2882, p= 0.7749). For a general overview of the behavior displayed by participants compared to the results in Doeller et al. (2008) see Supplemental Results.

**Figure 2:**
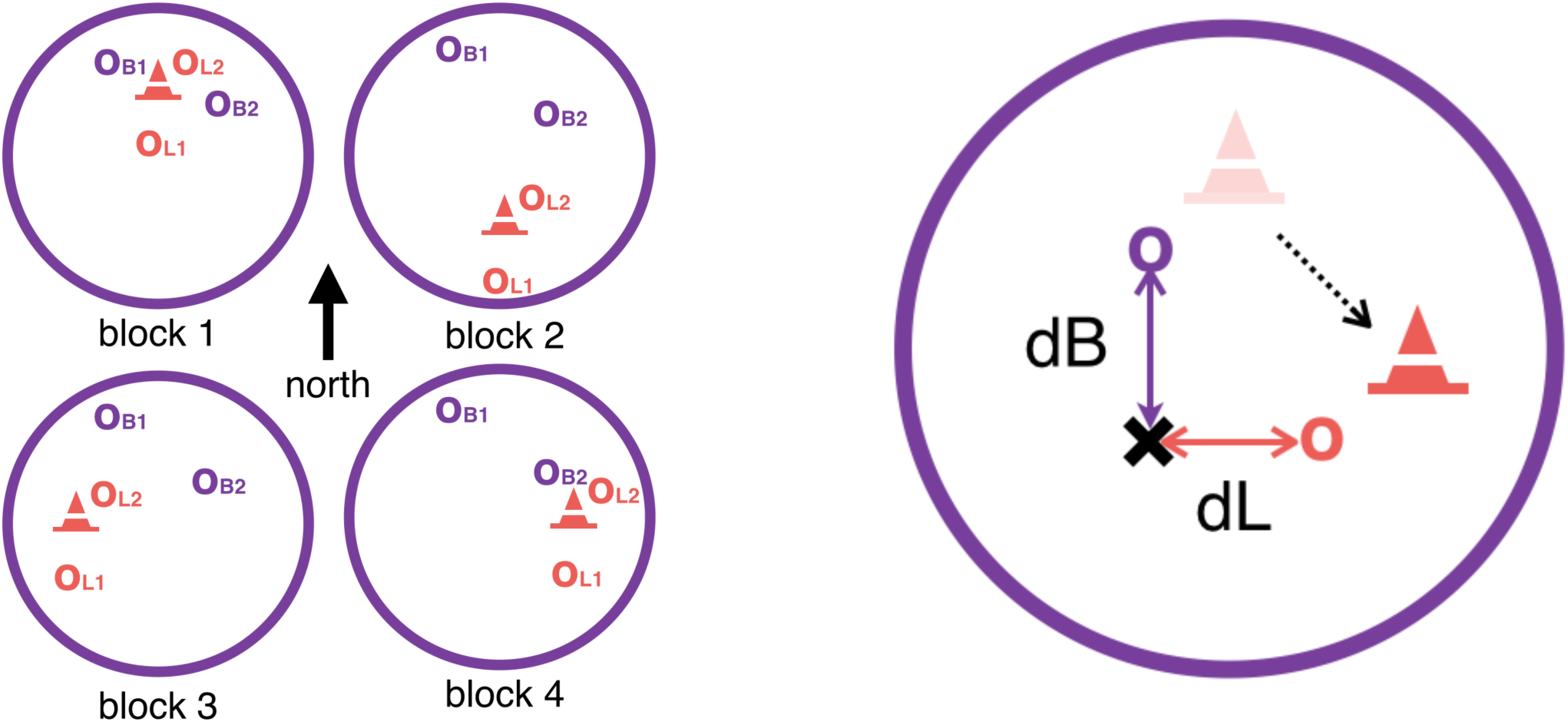
Left: Spatial Task Block Structure. Virtual arena as seen as from above. Between blocks the landmark (cone) moved in relation to the boundaries (large purple circle). Correct location of the two boundary objects (o_B1_ and o_B2_) stayed constant with respect to boundaries across all blocks. Correct location of the two landmark objects (o_L1_ and o_L2_) stayed constant with respect to the landmark across all blocks. **Right: Quantifying place and response memory.** The landmark (cone) moves (dotted line) in relation to boundaries (large purple circle) between blocks. In a trial before the landmark moves, an example object’s correct location (purple o), is in close proximity to the landmark (shaded cone). If participants remember this object location in relation to boundaries and distal cues, the predicted object location in the next block would also be indicated by the purple o. Conversely, if participants learned the object location in relation to the landmark,’ the predicted object location after the landmark moves to its new location (filled coned) would be the orange o. On the trials following movement of the landmark we thus operationalize place memory by the boundary distance error (dB) between their response (cross) and the location predicted by boundaries and distal cues (purple o). Response memory is operationalized by the landmark distance error (dL) between their response (cross) and the location predicted by the landmark cue (orange o). Lower dB and dL thus means better place memory and response memory, respectively.

We quantified place memory and response memory by, on the trials that followed the movement of the landmark, computing distance errors dB and dL (Doeller et al. 2008) relative to boundary and landmark cues (see Figure 2 Right). dB and dL measures are inversely related, respectively, to place memory and response memory.

We found a significant interaction effect of group (control vs. patient) by cue type (boundary vs. landmark) on distance error, while controlling for age and IQ, (*F*_1,97.58_=5.5080, *p*=0.021). This effect mainly reflected the finding that patient’s dB was significantly higher, i.e they displayed worse place memory (F_1,39.41_=2.5102, p=0.016) (see Figure 3 and Supplementary Table 1).

These analyses are collapsed over object type, that is, whether the correct location of an object was defined the boundaries or landmarks (participants were not instructed about this distinction); however, when additionally interacting group and cue type by object type we found no significant interaction (*F*_1, 780.8_=2.2040, *p*=0.138). These results are consistent with the prediction that anterior temporal lobe structures are necessary for place memory but not response memory.

**Figure 3:**
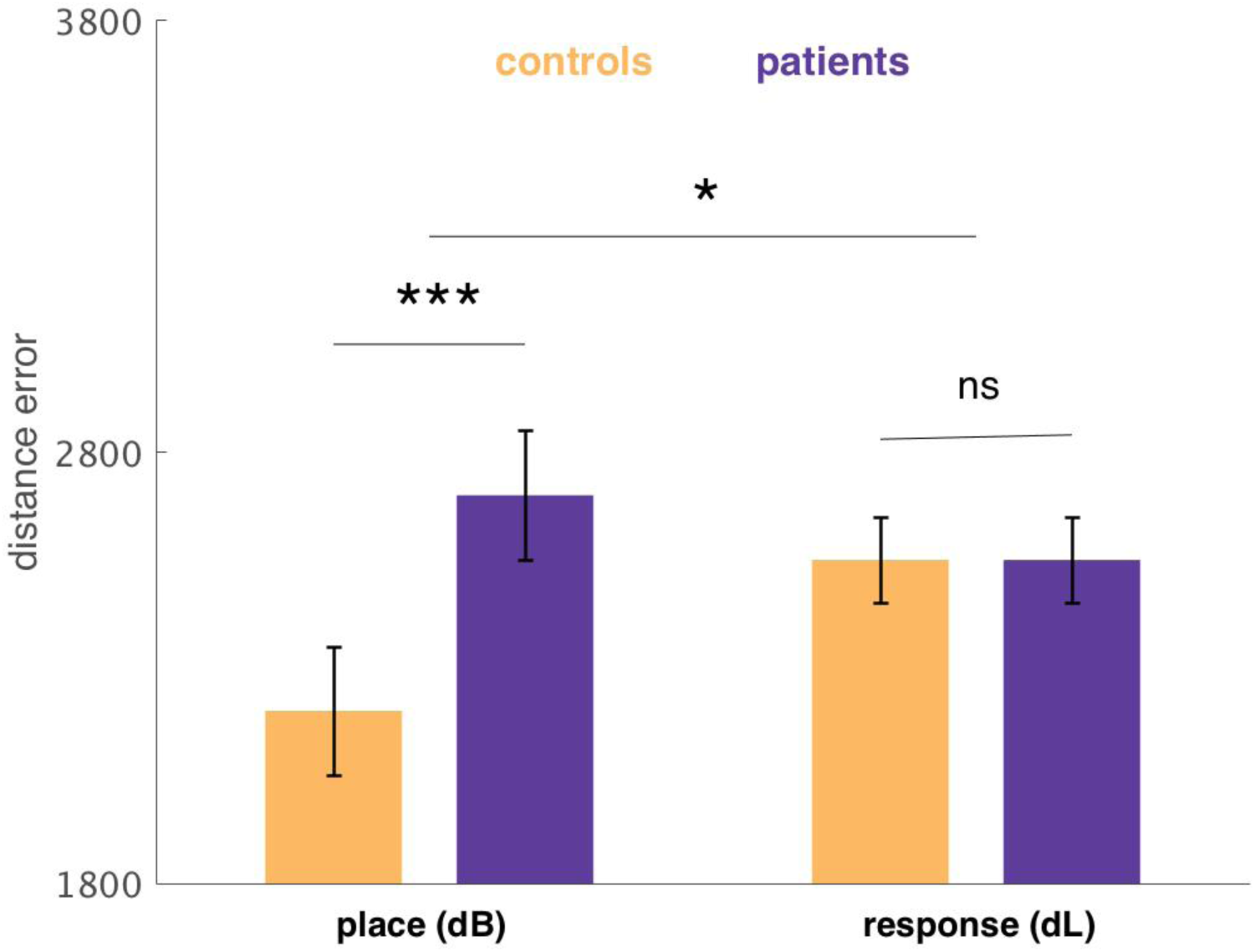
Average boundary (dB) and response (dL) distance error (arbitrary unit) for controls and patients. Estimated with a linear mixed-effects regression, controlling for IQ and age. Error bars indicate standard error. There was a significant group difference in boundary distance error (*F*_1,39.41_=2.5102, *p*=0.016), but not landmark distance error (*F*_1,85.3_=0, *p*=0.9990) with a significant interaction of group by cue type (*F*_1,97.58_=5.5080, *p*=0.0210).

### Patients display shift from model-based to model-free strategy

Participants completed 200 trials of a two-step Markov decision task (Daw et al., 2011) designed to quantify the reliance on model-based and model-free strategies (see Methods). On each trial the participant first made a choice between two spaceships. One spaceship most commonly (p=.7) transitioned to the purple planet, and otherwise made a rare transition (p=.3) to the red planet. For the other spaceship, probabilities were reversed. The participant then made a choice between two planet-specific aliens, each associated with a unique, slowly drifting probability of reward (see Figure 4). The mean number of completed trials was 195.3 (SE 2.1) with no difference between control and patient groups (t_38_=1.2515, p= 0.2188). The mean number of rewards received was 107.9 (SE 2.6), also, with no significant difference between control or patient group (t_36_= −0.0399, p= 0.9684). In general, rewards in this task are by design highly stochastic and not sensitive to differences in strategy.

**Figure 4:**
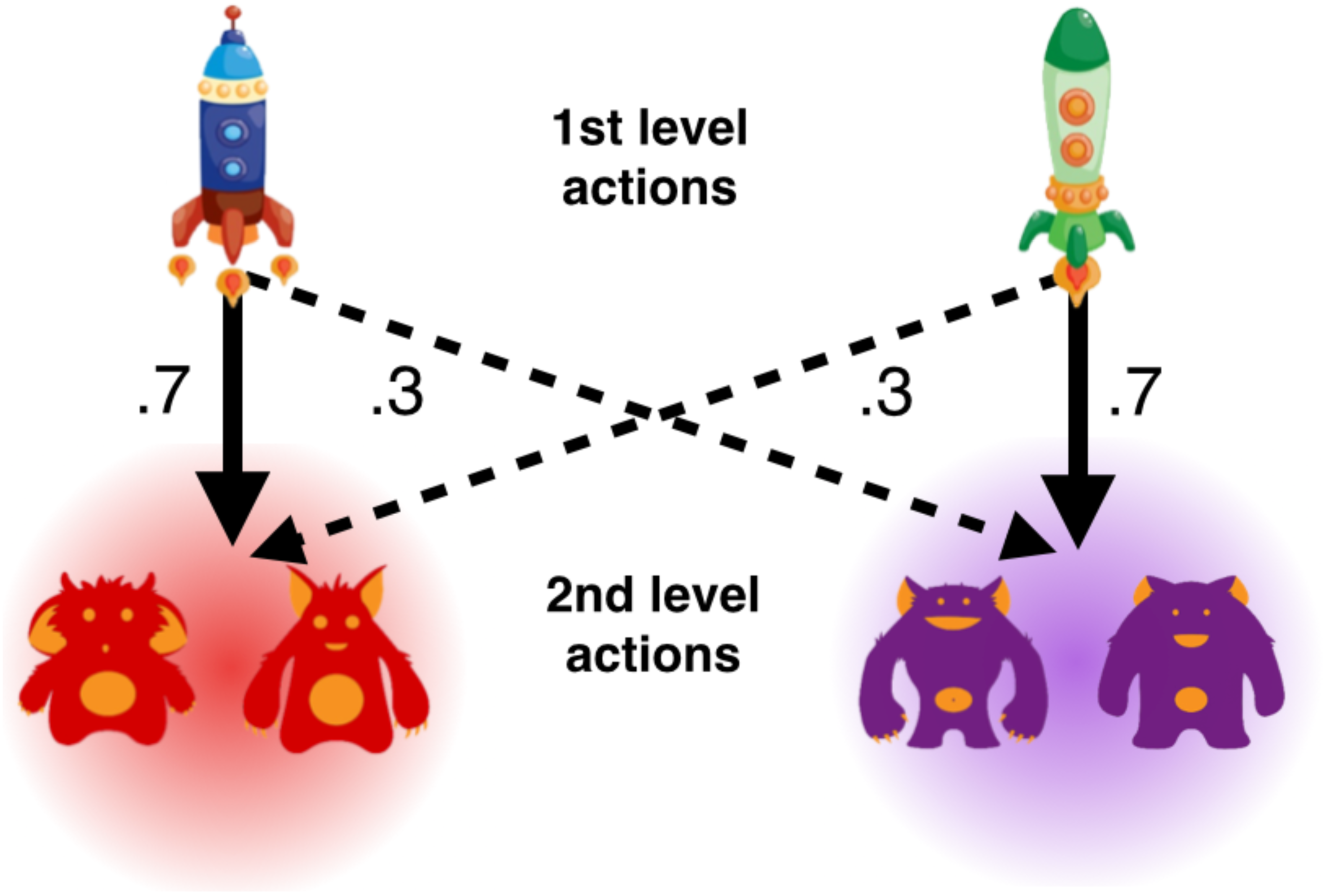
Two-step Markov Decision-Task. On each trial the participants chose one of two first level actions (spaceships). One space ship transitions the participant to red planet with *p*=. 7 while the other space ship transitions the participant to red planet with *p*=.3. Having transitioned to a second level state, participants chose between two second level actions (aliens) that were unique to each planet. Each alien was associated with a unique, slowly drifting, probability of receiving reward.

Figure 5 shows markers of both as a function of group, estimated from a factorial logistic regression (see Supplementary Table 2), which predicts choosing the same spaceship as on the previous trial. Model-free learning is signaled by a main effect of reward, i.e. a tendency to repeat choosing the spaceship that led to reward, whereas in model-based learning, choice of spaceship is mediated by expectations about the planets to which it leads, indicated by an interaction between reward and whether a rare or common transition occurred on the last trial. If, for instance, a reward is received but following a rare transition, a model-based agent should be less likely to repeat the choice of spaceship on the next trial. The difference in these effects measures the relative strength of model-based vs model-free choice.

**Figure 5:**
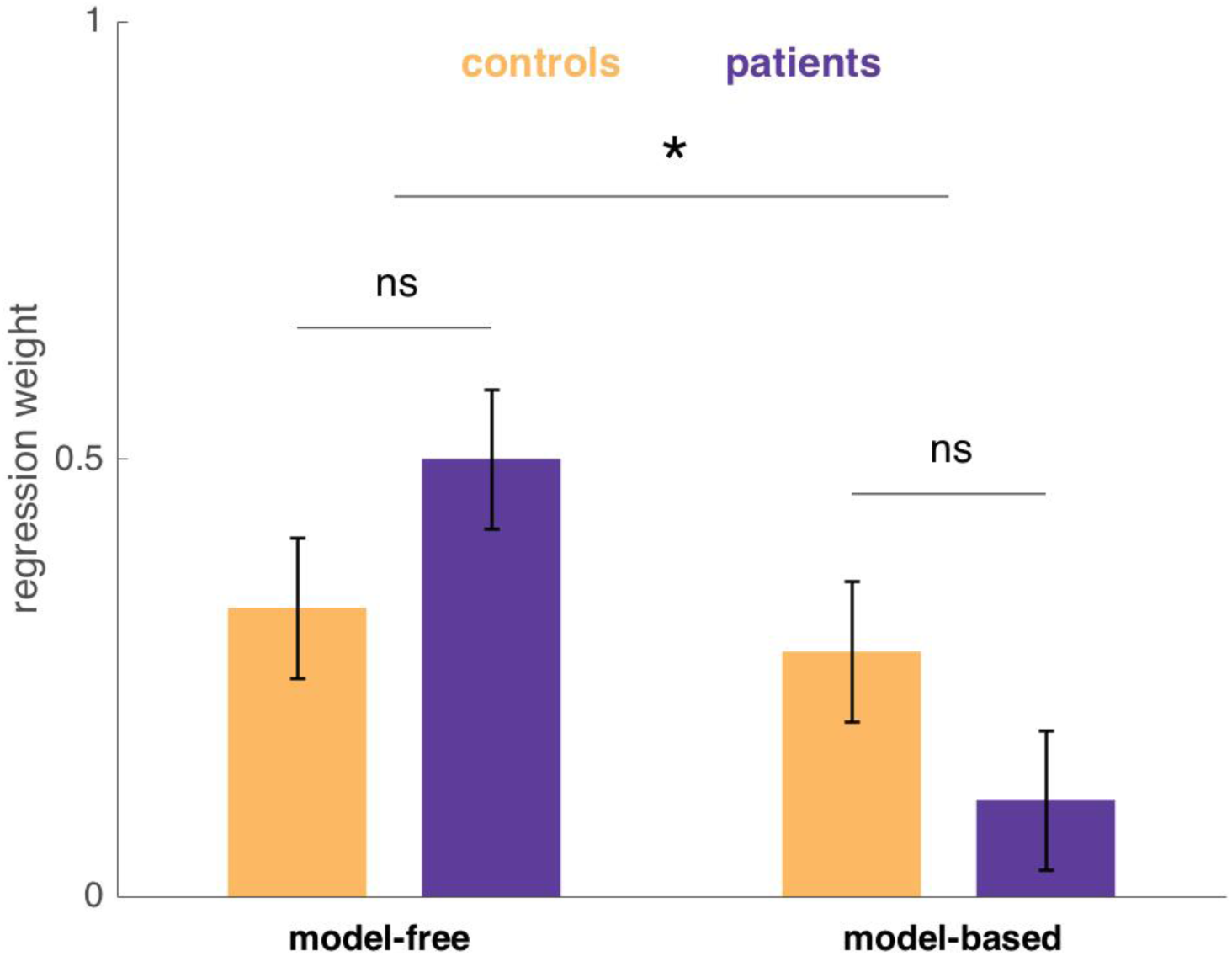
Model-free and model-based regression weights for controls and patients. Estimated with a logistic mixed-effects regression controlling for IQ and age. Error bars indicate standard error. The interaction of strategy (model-free vs. model-based) by group was significant (*z*=2.0278, *p*=0.043).

The expression of model-based vs model-free strategies differed significantly by group, with controls showing a relatively even mixture of strategies (similar to previous reports using this task) but patients being skewed away from model-based planning toward model-free learning (*z*=2.0278, *p*=0.043). This finding is consistent with the hypothesis that hippocampal damage in the ATL lesion group specifically affects the use of internal models or maps of task contingencies.

For simplicity, the above factorial analysis only considers the effect of the preceding trial’s events on each trial’s choice. To verify that our results were not dependent on this assumption, and in keeping with previous work (Daw et al., 2011) we repeated our analysis (see Supplementary Table 12) by fitting participants’ choices with a full 6-parameter computational learning model (Daw et al., 2011, as modified by Gillan et al., 2016), which uses the full history of preceding rewards to predict each choice. The results recapitulate the findings from the regression: chiefly, a significant interaction of RL-strategy and experimental group such that patients are biased away from model-based and towards model-free strategies (p=0.036). In addition, in this analysis (here going beyond the simpler regression analysis) the estimated strength of model-free learning is itself significantly higher in the patient group than the control group (p=0.043). The remaining parameters of the computational model did not differ significantly between groups.

### Model-based planning and place memory are correlated only in control group

So far, we have shown impaired spatial memory and model-based planning in the ATL group. Next, we compared the performance the two tasks, to examine their relationship and how that relationship is affected by ATL damage. If the hippocampus provides a common neural substrate for model-based planning and place memory, we should expect that these abilities be correlated in patients with intact hippocampi. Moreover, if this relationship depends on the intact hippocampus (or other ATL structures), the relationship should be correspondingly reduced in the lesion group. In other words, we wish to test the null hypothesis that any relationship between the tasks is unaffected by ATL damage, the rejection of which would support the alternative hypothesis that the ATL mediates such a relationship.

Since higher place memory performance is indicated by lower dB, for the following analysis, the participant-wise place memory performance will be expressed as the negative average boundary error (dB) for the participant, z-scored across all participants.

Figure 6 (also see Supplementary Table 3) displays the relationship between performances on the tasks, broken down by group. Place memory, as estimated from the spatial task, significantly predicted a participant’s use of model-based learning in the control group, even when correcting for IQ in the regression (*z*=2.6558, *p*=0.027). No such correlation was found in the patient group (*z*=0.1773, *p*=0.866). Indeed, the slope differed significantly between the two groups (*z*=2.1504, *p*=0.032); that is, there was a significant group difference in relationship between the two tasks’ measures of cognitive map usage. Furthermore, these effects appear, to be specific to model-based strategies, as there was no significant association between place memory and the degree of model-free learning in either group. Controlling for IQ, there was no overall effect of place memory on the model-free estimate in the control (z=0.1914, p=0.734) or patient group (z=1.2924, p=0.193), and no difference between groups (z=0.6856, p= 0.493). Finally, we repeated this analysis using the full computational learning model in place of the simpler regression-based index of learning (See Supplementary Table 13). Again, while controlling for IQ, we observed a strong positive correlation between model-based planning and place memory in the control group (p=0.04) but not the patient group (p= 0.847), although the group-wise interaction was not significant on this version of the analysis (p<0.1).

**Figure 6:**
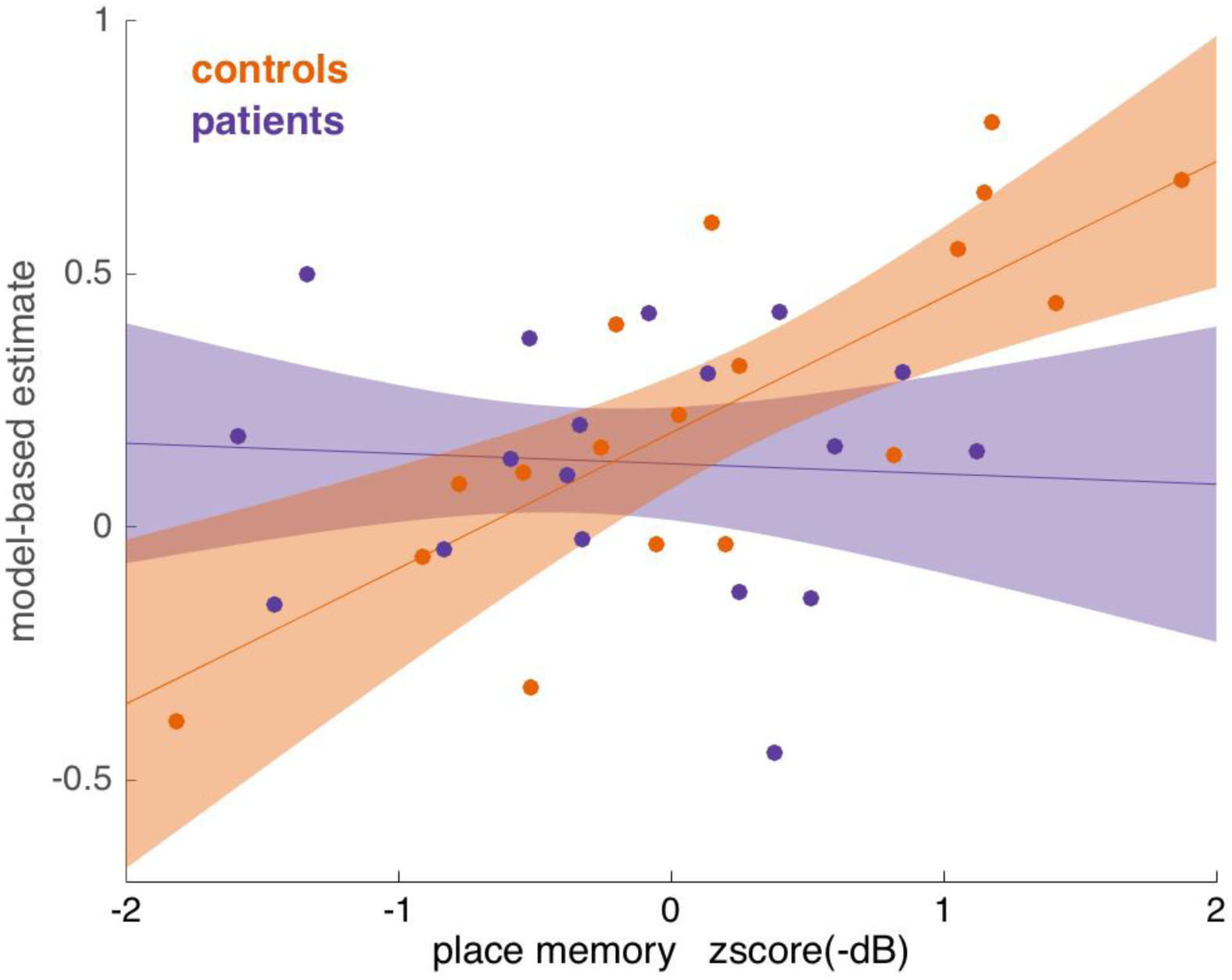
Relationship between model-based planning and place memory in controls and patients. Estimated with a logistic mixed-effects regression, controlling for IQ. Error bars indicate 80% confidence intervals. Individual place memory performances estimates are the negative average boundary error (dB), derived from the spatial task, z-scored across all participants. Dots indicate estimates for individual participants, calculated from the mixed-effects logistic regression. The trend was significant in the control group (*z*=2.6558, *p*=0.027), but not in the patient group (*z*=0.1773, *p*=0.866). The slope differed significantly between groups (*z*=2.1504, *p*=0.032).

### Deficits are more robust for patients with right lateralized ATL

Based on previous literature, we next sought to examine to what extent the reported effects might be preferentially associated with lesions lateralized to one side or the other. Breaking the data this way requires examining small subgroups (N=9 and 10), meaning that the key analyses comparing the two laterality groups against one another are underpowered relative to comparing either group to controls. Also, lesion laterality is correlated with overall lesion extent in our sample (Table 2). Altogether, these analyses are fundamentally more exploratory and their results more tentative than those reported above.

With those caveats, we expected place memory, and spatial relations generally, to be more strongly associated with the right hippocampus (Burgess et al., 2002). Accordingly, place-memory-related hippocampal activity in the fMRI study of the spatial memory task we used was right-lateralized (Doeller et al., 2008). It is less clear, a priori, how model-based planning should be lateralized.

Figure 7 Top and Bottom shows the spatial memory and decision task performance with the patient data further subdivided by ATL laterality; and Figure 8 shows the relationship between the two measures also broken down by laterality. The overall appearance in all three cases is that the differences between patients and controls (from Figures 4-6) were driven by the right lesion patients, with the left lesion patients more similar to controls. This impression is only partly borne out by statistics, however (see Supplementary Tables 4, 5 and 6). In particular, in all three cases the right patients differ significantly from controls (dB minus dL: F_1,92.22_=4.4635, p=0.034; model-based minus model-free: z=2.2950, p=0.022; across-task correlation between model-learning and place memory: z=2.5497, p=0.011), whereas the left patient group did not differ from controls in any case (dB minus dL: *F*_1,100.7_=2.644, *p*=0.107; model-based minus model-free: z=1.0008, p=0.317; across-task correlation between model-learning and place memory: z=0.4696, p=0.639). However, in no case were the lesion groups significantly different from one another (dB minus dL: *F*_1, 97.8_= 0.1503, *p*= 0.69903; model-based minus model-free: z=−1.1490, p=0.251; across-task correlation between model-learning and place memory: z=1.5524, p=0.121).

**Figure 7:**
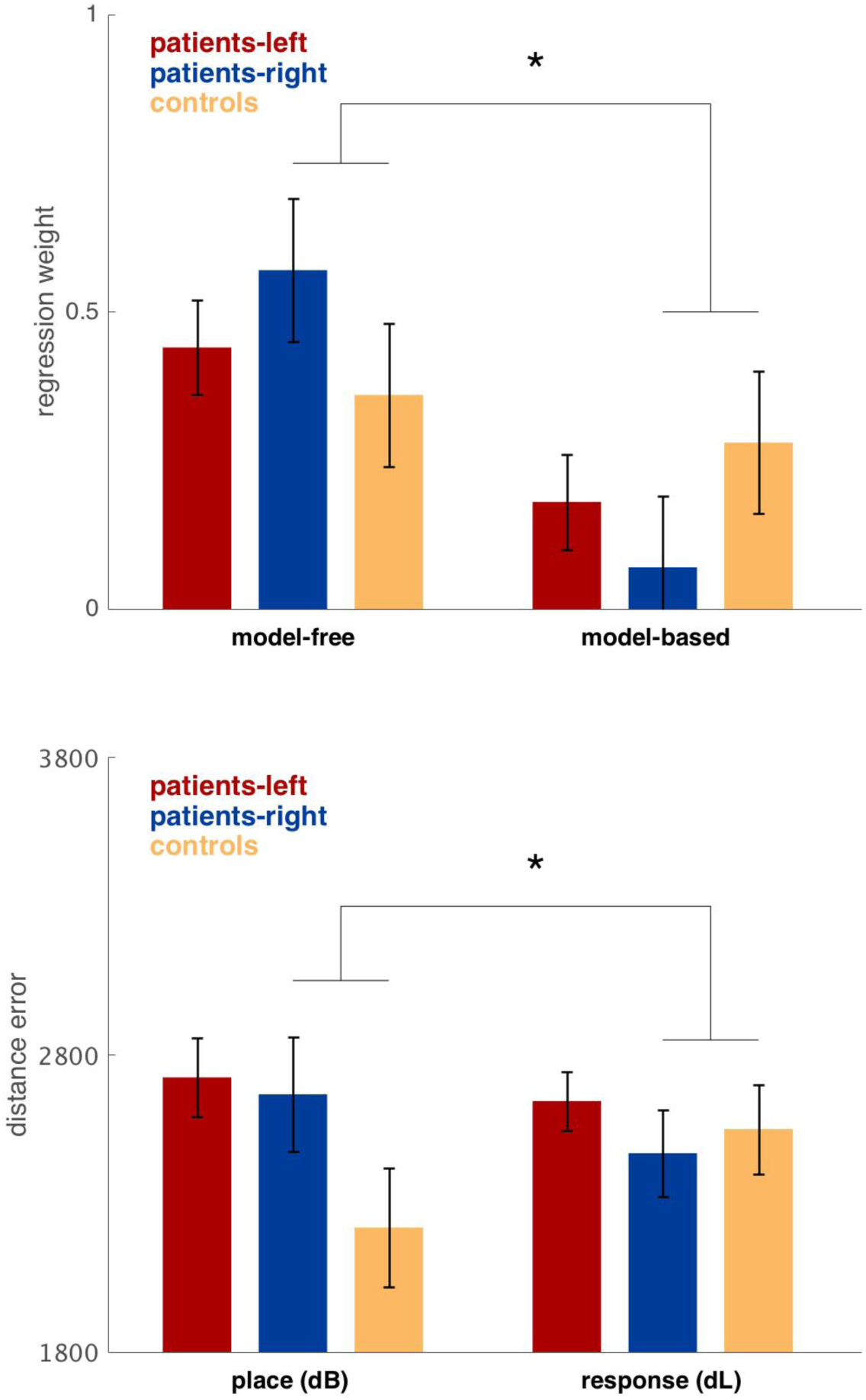
Top: Average boundary (dB) and response (dL) distance error (arbitrary units) for controls and patients with right and left lateralized ATL. Estimated with a linear mixed-effects regression, controlling for IQ and age. Error bars indicate standard error. There was a significant difference between dB and dL when comparing the control and right patient group (*F*_1,92.22_=4.4635 *p*=0.034). **Bottom: Model-free and model-based regression weights controls, right and left lateralized ATL patients.** Estimated with a logistic mixed-effects regression, controlling for IQ and age. Error bars indicate standard error. The difference in model-free vs. model-based was significantly different between the control and right patient group (*z*=2.295 *p*=0.022).

Thus, although noisy, there is a consistent suggestion across all three measures that the results of this study were most robust in the right lesion group.

**Figure 8:**
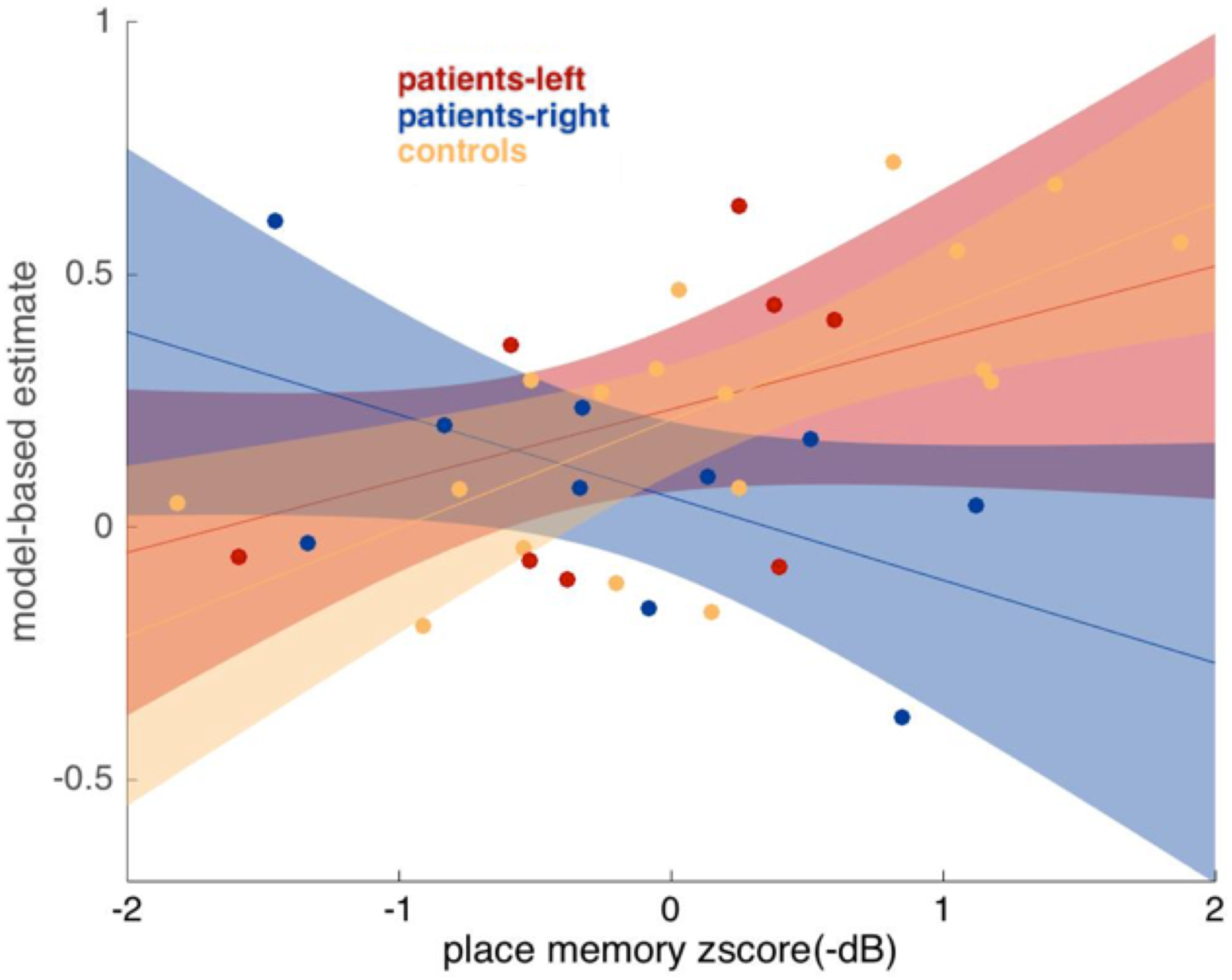
Relationship between model-based planning and place memory performance for controls and patients with right and left lateralized ATL. Estimated with a logistic mixed-effects regression, controlling for IQ. Error bars indicate 80% confidence intervals. Individual place memory performances estimates are the negative average boundary error (dB), that were derived from the spatial task, z-scored across all participants. Dots indicate estimates for individual participants, calculated from the mixed-effects logistic regression. The trend differed significantly between the control and right patient group (z=2.5497, p=0.011).

### Size of lesion to right hippocampus predicts planning deficits

One way to sharpen the foregoing analyses is to focus specifically on not just the side but the particular anatomical region hypothesized to underlie the effects: the hippocampus. Accordingly, we tested how performance on the tasks co-varied with estimated lesion size on the right and left hippocampus respectively. Hippocampal lesion size for each patient (see Figure 1) was estimated by comparing the normalized anatomical masks to the Harvard-Oxford Lexicon (p>.5) (see Methods). Importantly, the ATL procedure involves a pattern of damage to numerous temporal lobe structures in addition to hippocampus, which means one should be highly cautious interpreting results with respect to any particular structure. Although it is not practical to control for damage to many different MTL structures individually, we attempt to mitigate these concerns and focus on hippocampal lesion size by controlling for the overall lesion size as a nuisance effect. The regression analyses also controlled for age and IQ.

Figure 9 Top and Middle panels (see Supplementary Table 7 and 8 respectively) show the effect of hippocampal lesion size, of either laterality, on place memory (negative z-scored boundary error dB) and model-based planning. Model-based planning was significantly worse for larger right hippocampal lesions (*z=2.8157, p*=0.005). Conversely, planning was not significantly related to the amount of hippocampal damage on the left hippocampus (*z=0.1100, p*=0.9124) and the difference between right and left effects was significant (*z*=2.4702, *p*=0.0140). For place memory, the data trended in a similar direction with impairments being related size of lesions in the right (*F*_1,12.16_=2.6082, 0.1320) but not the left hippocampus (*F*_1,12.93_=0.1204, *p*=0.7340). These results further suggest that in addition to being critical to place memory, the right hippocampus facilitates model-based planning.

**Figure 9:**
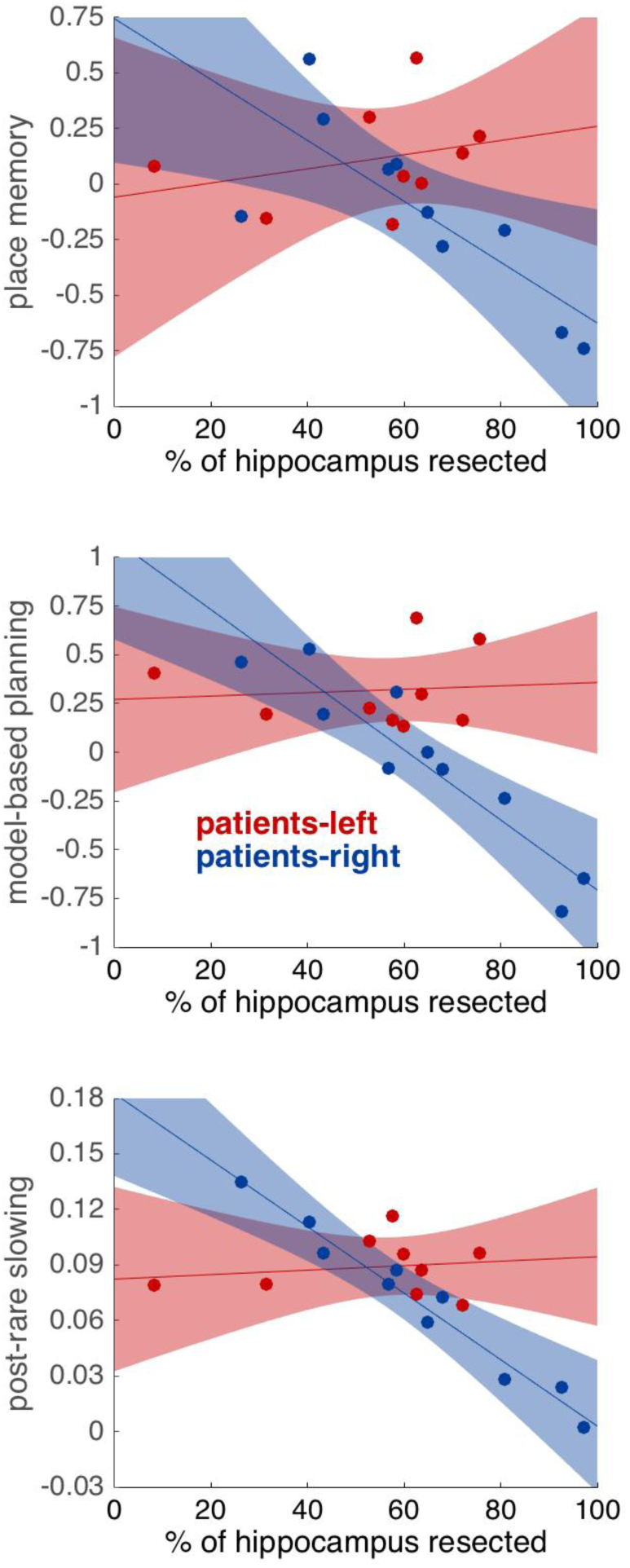
Relationships between variables of interests and estimated percentage of hippocampus resected, for patients with right and left lateralized ATL. Estimated with a mixed-effects regressions, controlling for IQ, age, and total size of lesion. Error bars indicate 80% confidence intervals. Dots indicate estimates of effects for each individual participant, derived from the mixed-effects regression. **Top: Relationship between place memory (negative z-scored dB) and hippocampal lesion size.** There was a trending relationship in the right patient group (*F*_1,12.16_=2.6082, 0.1320) but not in the left patient group (*F*_1,12.93_=0.1204, *p*=0.7340) with a trending difference between the right and left patient groups (*F*_1,12.27_=2.5889, *p*=0.1330). **Middle: Relationship between model-based planning (regression weight from logistic regression) and hippocampal lesion size. The trend was significant in the right patient group (*z*=2.8157, *p*=0.005) but not in the left patient group (*z*=0.1100**, *p*=0.9124), with a significant difference between the right and left patient groups (*z*=2.4702, *p*=0.0140). **Bottom: Relationship between post-rare slowing (2^nd^ level reaction time difference following rare transitions vs. common transition) and hippocampal lesion size.** The trend was significant in the right patient group (*F*_1,18.7_=9.9930, *p*=0.005) but not in the left patient group, with a significant difference between the right and left patient groups (*F*_1,18.7_=7.787, *p*=0.012).

### Size of right hippocampal lesion predicts loss of knowledge of task transition structure

Finally, we attempted to further elucidate what aspect of model-based planning was supported by right hippocampus. If the hippocampus supports knowledge of the task transition structure, a necessary antecedent for model-based planning, hippocampal damage should affect other measures of knowledge of that transition structure. In particular, in the two-step decision task, people respond more slowly for the second-level choice (aliens) when the state they arrived at followed a rare rather than a common transition (Deserno et al., 2015). This presumably reflects less preparation of a response for the expected state (Dezfouli and Balleine, 2013). Such post-rare slowing implies knowledge of the transition structure, but importantly, it is dissociable from model-based planning, in that post-rare slowing is observed even in participants (such as children; Decker et al., 2016) who do not show significant model-based planning.

In the present data, model-based planning and post-rare slowing were indeed correlated across participants (z=3.5327, p<0.001) (see Supplementary Table 9), with no differences in this correlation between any of the groups, and individually significant in each group (with the exception of the right patient group where the correlation only reached trend level z=1.6979, p=0.09) (see Supplementary Table 10). Thus, to the extent that any participant exhibits model-based choices, this effect is closely related to post-rare slowing.

While there were no significant differences between groups in reaction time following rare compared to common transitions, when relating the post-rare slowing effect to lesion size (see Supplementary Table 11) we observed a selective effect of lesion size in the right hippocampus (*F*_1,18.7_=9.9930, *p*=0.005) which was significantly different from left hippocampus, (*F*_1,18.7_=7.787, *p*=0.012); such that larger right hippocampal lesions reduced the post-rare slowing effect (see Figure 9 Bottom). This is consistent with the hypothesis that the right hippocampus subserves knowledge of the task transition structure, which is necessary for planning.

## Discussion

Although extensive evidence indicates that the hippocampus supports localization in allocentric space, there is relatively little direct evidence for the hypothesis that the same mechanisms extend to model-based planning. We addressed this gap by testing model-based planning and place memory in patients with extensive hippocampal damage as a result of unilateral ATL lesions and matched, neurologically typical controls. Our results are consistent with the hypothesis that both place memory and model-based planning share a common mechanism, which is affected in ATL patients and, more tentatively, associated with right hippocampus.

As predicted, ATL patients displayed significantly attenuated place memory in our MWM-like spatial memory task, alongside spared response memory. These results corroborate the dual-systems view of navigation supported both by rodent lesion (O'Keefe and Nadel, 1978, McDonald and White, 1994, Packard and McGaugh, 1996, Pearce et al., 1998) and human neuroimaging experiments (Hartley et al., 2003, Iaria et al., 2003, Voermans et al., 2004, Doeller et al., 2008), whereby the hippocampus supports fast learning of allocentric spatial maps and the striatum facilitates slow, incremental associations between stimuli and responses.

The patients were also significantly biased away from using model-based planning and toward model-free habitual strategies in the two-step Markov decision task. This result provides causal evidence for the inference that temporal lobe structures support model-based planning, over and above their role in place memory. The appearance of a compensatory shift toward improved model-free learning, which is rarely reported with this task (Frank et al., 2004), indicates that behavior in the ATL patients is not simply noisier, and instead is consistent with models invoking multiple, potentially competing, reinforcement-learning systems in the human brain (Daw et al., 2005). Still, our results do not speak clearly to the perennial question whether the hippocampus plays a special role in such models for spatial vs. more abstract relational tasks. This is because although our planning task is structured like an abstract Markov decision process, its cover story, in terms of rocket trips to planets, might have elicited a spatial interpretation.

Our results also complement and extend previous research with rodents. Unit recording studies have shown results suggestive of hippocampal involvement in model-based planning, notably replay of forward trajectories in hippocampal place cells (Johnson and Redish, 2007, Pfeiffer and Foster, 2013). However, in contrast to the current study, studies with place cell recordings have not yet shown behavioral evidence for a link between the hippocampus and the use of this knowledge in planning, nor do they provide evidence for a causal role of hippocampus in such a function. In these respects, our results more closely parallel a recent report of a related deficit in model-based learning in rodents during inactivations of the dorsal hippocampus, using an analogous two-stage Markov decision task (Miller et al., 2017). The targeting of the inactivation to hippocampus in the rodent study sharpens the anatomical specificity of the effect. Conversely, our human result clarifies the contribution of the damaged structure, because we know more about the computations underlying model-based behavior on this task in humans. In particular, in humans, but not yet rodents, model-based choices have been explicitly linked to prospective neural activity at decision time (Doll et al., 2015). This helps to rule out other potentially confounding strategies, such as that the apparently model-based choices in rodents are instead produced by some learned response switching rule contingent on events spanning multiple trials. It has been suggested that such model-free strategies might arise following overtraining of the sort used to teach animals this task (Akam et al., 2015, Economides et al., 2015); this might also implicate hippocampus for other, confounding reasons, such as its involvement in trace conditioning and latent states (Solomon et al., 1986, Büchel et al., 1999, Gershman et al., 2010). Our findings of a similar result in humans, without extensive training, thus help corroborate the interpretation of the rodent study as well.

Comparing performance between our two tasks we also found that in healthy controls, place memory performance correlated with the extent of reliance on model-based planning strategies. Importantly, no such correlation was found in the patient group. These results suggest that some common substrate for the tasks is provided by the temporal lobes in the healthy brain. Following damage to this structure, however, the two tasks may be supported by differential compensatory mechanisms, leading to their decorrelation. Altogether, these findings are consistent with the hypothesis that both model-based planning and place memory share a common mechanism, which is impaired in ATL patients.

It is surprising and interesting that the effects we report emerge following damage to only one lobe, as unilateral temporal lobe damage is generally known to produce rather subtle effects on cognition in humans (Spiers et al., 2001) and animals (van Praag et al., 1998), relative to the famously dramatic effects of bilateral lesion (e.g. Scoville and Milner, 1957). This may relate to our use of two behavioral tasks that are well attuned to temporal lobe function. However, there exist inherent and important caveats in drawing conclusions about the neural bases of effects from a study of this sort. It is possible that the observed effects are caused by damage to the brain, including the hemisphere not surgically altered, as a result of the chronic epilepsy that prompted the surgery. Indeed, as with all studies of temporal lobe function in patients with epilepsy, the possibility that impaired behavior and cognition in patients is due to a history of epilepsy rather than the surgical intervention per se must be taken into account.

For this reason and others, we must also be cautious about associating the damage with individual structures. Our analyses indicate that the size of lesion to the right hippocampus is significantly related to model-based planning deficits and has a trending relationship to place memory deficits. Still, ATL lesions additionally affect a number of other regions including parahippocampal cortex, perirhinal cortex, and amygdala, which might also subserve these effects. Moreover, since the pattern of the lesions mainly varies in the extent that the temporal lobe has been removed in the dorsal direction, the patterns of damage to these structures tends to covary across individuals. Such collinearity makes it difficult to use variation across patients in damage to individual structures to fully disentangle their differential roles. We attempted to mitigate these issues by controlling for overall lesion size. Nevertheless, due to the very substantial analytic and interpretational issues, this anatomical specificity remains emphatically tentative.

These caveats aside, a final question posed by our results concerns how place memory and model-based planning actually relate to one another. In the spatial literature, the notion of a cognitive map primarily refers to place-selective hippocampal activity, which allows organisms to recognize and remember discrete locations. From the perspective of planning, a cognitive map goes beyond such a representation, but is built upon it: the map captures the relationships between locations, which can be used to evaluate candidate actions. This function fits well with the broader view of hippocampus supporting relational memory (Eichenbaum and Cohen, 2001, Davachi and Wagner, 2002, Schapiro et al., 2016, Boorman et al., 2016, Garvert et al., 2017) and with our analyses of reaction times, which indicate that the planning deficit in patients stems from hippocampal damage being accompanied by attenuation of the knowledge of relationships between actions and states. This view is also consistent with recent computational models describing how the hippocampus might serve model-based planning. In spatial tasks, sequential activations of place-selective cells are hypothesized to provide, not only a mnemonic function through supporting reactivation of previously traversed trajectories, but a planning function by generating novel place cell sequences, based on the learned contingencies between locations (Johnson and Redish, 2007, Pfeiffer and Foster, 2013, Mattar and Daw, 2017). The related successor representation model (Stachenfeld et al., 2017, Garvert et al., 2017) also focuses on learned relationships among locations, by proposing that place selectivity itself is built from experience of state transitions to reflect expectations about future locations. A key challenge for future work addressing these ideas will be studying hippocampal activity in tasks, like the planning one used here, which manipulate animals' experience of such environmental relationships and reveal how they leverage this knowledge to guide their choices.

## Acknowledgements

We wish to thank Philip Parnamets and Aaron Bornstein for helpful input. This project was supported by NIH grant DA038891, part of the CRCNS program, and a gift from Google DeepMind.

## Methods

### Participants

The patient sample consisted of 19 individuals who had undergone unilateral anterior temporal lobectomy (ATL) for the treatment of intractable epilepsy. Patients were recruited from the New York University (NYU) Patient Registry for the Study of Perception, Emotion and Cognition (PROSPEC). The control sample consisted of 19 healthy controls (HC), who were recruited from the local community through internet-based advertisement and gave consent to participate in the study.

A clinical neuropsychologist (MRM or KB) conducted all standardized procedures for screening patients for inclusion into NYU PROSPEC. Patients were only selected for inclusion if there was no evidence of global cognitive dysfunction as measured by a comprehensive neuropsychological evaluation, an FSIQ (Wechsler Adult Intelligence Scale-Fourth Edition (Wechsler, 2008) above 80, no evidence of diffuse atrophy on MRI (e.g., brain tumor or idiopathic epilepsy), or and no history of psychiatric or neurologic disease other than the primary etiology for the focal brain lesion.

On the day of testing, participants completed the two behavioral tasks separated by a short break. For all participants the sequential decision making task was given first, followed by the spatial memory task. For the control participants the completion of the tasks was followed by the administration of the WAIS-IV. For the patient participants, the WAIS-IV had been completed during screening procedures for inclusion into PROSPEC.

### MRI Scanning and Image Processing

When post-surgical structural brain scans (T1 MP-RAGE) were not available from the referring center, the Department of Radiology at the NYU School of Medicine, patients were imaged at the NYU Center for Brain Imaging on a 3-Tesla Siemens Allegra head-only MR scanner. Medical Center scans were obtained using 1.5 or 3-Tesla Siemens full-body MR scanners. Image acquisitions included a conventional three-plane localizer and two T1-weighted gradient-echo sequence (MP-RAGE) volumes (TE = 3.25 ms, TR = 2530 ms, TI = 1.100 ms, flip angle=7°, FOV=256 mm, voxel size=1×1×1.33 mm). Acquisition parameters were optimized for increased gray/white matter image contrast.

The high-resolution structural images from each patient were normalized to Montreal Neurological Institute (MNI) standard space using FSL FLIRT (FMRIB’s Linear Image Registration Tool; http://fsl.fmrib.ox.ac.uk/fsl) (Jenkinson and Smith, 2001). This consisted of a two-step procedure: First, using MRIcron (http://www.mccauslandcenter.sc.edu/mricro/mricron/), a mask was drawn over the lesion and any craniotomy defect to prevent bias in the transformation, then masked voxels were assigned a weight of “0” and ignored during a subsequent 12-parameter affine transformation of the lesioned brain to the standard MNI 1 mm reference volume (Mackey et al., 2016). The second step was manually tracing the lesions on individual slices of the patients’ brains overlaid on the standard MNI brain template, while crosschecking in all three planes. This tracing procedure produced a 3D mask with “1” indicating the presence of the lesion and “0” the presence of normal tissue. All patients had surgical lesions, which made the margins readily visible on the T1-weighted MRI images. In instances where there was uncertainty regarding the lesion margins, the treating neurosurgeon(s) and/or neuroradiologists were consulted.

Lesion masks drawn in MNI space were subsequently overlaid on the Harvard-Oxford Structural Atlas) (Mazziotta et al., 2001) to estimate the extent of damage to the hippocampus (see Figure 1). Since the Atlas was also defined in MNI space, hippocampal lesion size was calculated as the voxel overlap between the individual lesion masks and the hippocampus as defined by the Atlas with p>.5.

### Spatial Memory Task

Each participant completed 64 trials of a spatial memory task, identical (with one exception, see below) to the task used by Doeller et al. (2008). On each trial, participants navigated a virtual reality arena using keyboard presses. UnrealEngine2 Runtime software (Epic Games) was used to present a first-person perspective view of the arena. The virtual arena was bounded by a circular wall, contained a single intra-maze landmark in the form of a traffic cone, and was surrounded by distant cues (mountains, clouds, and the sun) projected at infinity. Both the boundary (wall) and landmark (cone) were rotationally symmetric, leaving the distal cues as the main source of orientation.

At the beginning of each trial, a picture of one of four objects was presented on a grey background for 2 s. Participants were then placed in a random position within the arena without any objects, one-fifth of the radius from the center of the arena and facing a random direction (note that in Doeller et al. (2008) the starting radius was not restricted). Participants subsequently had 12 seconds to navigate to the correct location of the object as they remembered it from previous trials, and indicate that position by a button press. Following this button press, the object immediately appeared in its correct location. If no response had been made in 12 seconds, the object also appeared in its correct location automatically. Participants ended the trial by collecting the object in its correct location. A fixation cross was then presented for 2 s, before the start of the next trial.

The task consisted of 64 trials divided into 4 continuous blocks, each containing 4 pseudo-randomized presentations of each of the 4 objects. Between blocks, the landmark moved in relation to the boundaries, such that there were four arena configurations, with the landmark roughly in the middle of the north, south, west, and east sectors of the arena, as defined by the distal cues (See Figure 2 Left). The order of arena configurations over blocks was counterbalanced across participants and experimental groups. Participants were not informed of the landmark movements prior to the experiment.

During the first block, the correct location of all objects was in rough proximity of the landmark. Two of the objects were ‘boundary objects’, for which the correct locations were fixed relative to the environmental boundaries across the whole experiment. The other two objects (unannounced to the participants) were ‘landmark objects,’ for which correct locations were fixed at a constant distance and direction to the intra-maze landmark even as the landmark moved.

The task probed for memory of correct object locations within the arena. Critically, by manipulating the landmark location in relation to the boundary and distal cues, the task distinguished whether participants stored place memory of allocentric location in relation to the boundary and distal cues, or by egocentric response memory in relation to individual landmarks (See Figure 2 Right). The original study, using the same procedure during fMRI in healthy participants, showed that place and response memory correlated with activity in the right posterior hippocampus and striatum, respectively (Doeller et al 2008).

Participants practiced in an unrelated virtual environment with a different set of object stimuli before performing the experiment. Additionally, before the first trial, participants collected each of the objects once in their correct block 1 locations.

### Spatial Memory Task Analysis

To measure place and response memory we focused our main analysis on the trials following the relative movement of the landmark in relation to the boundaries. Place memory was quantified by boundary distance error (dB), where dB was the distance from the response location to the correct location as defined by the boundary and distal cues in the previous block (See Figure 2 Right). Response memory was quantified by landmark distance error (dL), where dL was the distance from the response location to the correct location as defined by the landmark, according to the position of the object relative to the landmark in the previous block, translated with respect to the landmark’s new position (See Figure 2 Right). Low dB thus indicated better place memory, and low dL better response memory. While it is true that knowing the correct location in relation to the landmark also requires orientation from the landmark and thus relies on distal cues, it has been shown that rats with hippocampal lesion are not impaired at learning location this way (Pearce et al., 1998).

To capture the repeated-measure structure of the data, all statistical analyses of performance in the task were done using mixed-effects linear regression. The models were estimated using the *fitlme* function in Matlab, with standard errors computed using the Satterthwaite approximation to the degrees of freedom when the model was linear, and Wald (asymptotic Gaussian) test for logistic models. The dependent variable, distance error (dB and dL respectively for each trial) was regressed on the key explanatory variables lesion group, error-type (boundary or landmark) and object type (boundary or landmark), while also controlling for additional nuisance explanatory factors, age and IQ.

### Sequential-Decision Making Task

Participants also completed 200 trials of a two-step Markov decision task designed to quantify the extent to which participants use a world model to prospectively evaluate actions (Daw et al., 2011). The task was framed as a game about mining for space treasure (Decker et al., 2016). Each trial involved two choices in succession, followed by reward (See Figure 4). Participants first made a choice between two actions, depicted as spaceships, randomly presented on the left and right. The choice resulted in a transition to one of two second-stage states (depicted as a red or purple planet). One spaceship most commonly (p=.7) transitioned to the purple planet, and otherwise made a rare transition (p=.3) to the red planet. For the other spaceship, probabilities were reversed. Participants were informed that each spaceship was more likely to go to a different planet but not which planet, nor the explicit transition probabilities.

Subsequently, participants made a choice between two actions depicted as a pair of aliens that were unique to the planet, randomly presented on the left and right. Each alien was associated with a probability of monetary reward (vs nothing) that slowly diffused over trials according to an independent random walk. Rewards were paid out at the end of the experiment at a rate of 15 cents per reward. The random change in the second-stage reward probabilities encouraged participants to adjust their choice preferences at both stages trial-by-trial, so as to maximize payoffs. For each choice, participants had 3 seconds to respond; or else the trial was aborted with a time-out message.

Prior to the experiment, participants completed an extensive instructional tutorial. The tutorial included a 20-trial practice run, using a different set of visual stimuli (planets, spaceships, and aliens) but otherwise identical.

### Sequential-Decision Making Task Analysis

The logic of the task exploits the noisy coupling between spaceships and planets to measure model-free learning - directly learning the value of spaceship choices vs. model-based planning - prospectively computing the value of the spaceship choices in terms of the planets they lead to.

For instance, consider on some trial choosing the spaceship that usually transitions to the purple planet, but instead being taken to the red planet (a “rare” transition). On the red planet your choice of alien is subsequently rewarded. In this situation model-free and model-based strategies make conflicting predictions about first-level choice behavior on the next trial. Participants using a model-free strategy will be more likely to choose the same spaceship on next trial, as it was rewarded. Conversely, participants using a model-based strategy will be more likely to switch and choose the other first-level action. This is because the model-based strategy computes the value of the spaceships using a cognitive map or model of their transition probabilities to the respective planets and the reward expected at the planets.

The goal of analysis was to estimate, for each participant, the extent to which they followed either strategy. Following previous work (Daw et al., 2011), we did this two ways, using a factorial logistic regression that captures the above qualitative logic, and fits of a more elaborate, but more assumption-laden, computational learning model.

We analyzed the first-level choices over spaceships using mixed-effects logistic regression (estimated using the *fitglme* function in Matlab). For each trial, the dependent variable (coded as stay with the same spaceship or switch, relative to the previous trial) was explained in terms of two events from the previous trial: whether reward was received, whether the planet encountered was reached following a common or rare transition given the spaceship chosen, and the interaction of these two factors. Our measure of model-free choice was the main effect of reward; our measure of model-based choice was the interaction of reward by transition type (common vs. rare). We further interacted the task factors with experimental group (lesion vs. control) as well as with two nuisance covariates, IQ and age, which have both been shown to affect behavior on this task (Gillan et al., 2016). The intercept, and the regression coefficients for reward, transition, and their interaction were all taken as random effects (allowed to vary across participants).

To test our predictions about the relationship between reinforcement learning strategies employed in the decision-making task and place memory performance from the spatial memory task, we also specified a second regression model which interacted the task- and group-related factors (reward, common vs rare, group, and their interactions) with participant-specific z-scored estimates of place memory (dB) derived from the spatial memory task. (IQ was again included as a nuisance variable.) The interactions with dB (up to four-way) measure the extent to which the various effects in the decision task systematically vary, across participants, with their spatial memory performance; i.e. this is analogous to extracting per-participant effect sizes from the decision model and correlating them with dB, but by estimating that correlation as an effect within the regressing defining those decision effects, accounts properly for uncertainty in the per-participant estimates.

Individual dB measures were extracted as the random effects from a mixed-effects linear regression, in which the dependent variable was per-trial dB and the intercept was taken as a random effect across participants. No explanatory variables except for the intercept were included in this model. This is analogous to defining dB as the average per-trial error for each participant, but regularizes these estimates over the whole population of participants.

## Supplementary Methods for Computational Model Fit

The logistic regression analysis considers only the previous trial’s experience in predicting each choice; this simplification is motivated by a limiting argument over the learning rate parameter in a more elaborate RL model of the data (Daw et al., 2011). In order to ensure that our results were not affected by neglecting the effect of earlier trials, we repeated our analyses fitting each participant’s trial-by-trial choices with a full RL model in which each choice depends on values learned from all previous rewards (based on Daw et al., 2011, but using the version from Gillan et al. 2016). To estimate the model we utilized Markov Chain Monte Carlo (MCMC) methods, implemented in the Stan modeling language (Stan Development Team). Given an arbitrary generative model for data dependent on free parameters, the method permits samples to be drawn from the posterior probability distribution of parameter values, conditional on the observed data. From the quantiles of these distributions, we constructed confidence intervals – technically, credible intervals – over the likely values of the free parameters (Kruschke, 2010). We also report the posterior likelihood that the credible region contains zero, as one minus the size of the largest symmetric credible interval that excludes zero, which is roughly comparable to a two-sided P value.

For each model, we produced 4 chains of 10,000 samples each. The first 250 samples from each chain were discarded to allow for equilibration. We verified the convergence of the chains by visual inspection, and additionally by computing for each parameter the ‘potential scale reduction factor’ 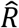 (Gelman and Rubin, 1992). For all parameters, we verified that 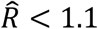, a range consistent with convergence (Gelman, Carlin, Stern, and Rubin, 2003).

We simultaneously estimated a model of all the data, incorporating individual parameters for each participant nested within a population-level model of the distribution of these parameters for each group.

At the participant level, the model is the same as the one used by Gillan et al. (2016), and full equations are presented there. In brief, the model learns from experience to predict values *Q*(*s, a*) for the different actions *a* (rockets, aliens) in the different states (planets and the starting state). Different RL algorithms, model-based and model-free, produce different estimates *Q* at each step. First-level (spaceship) choices are determined by softmax choice, according to the weighted combination of model-based and model-free *Q* values, with weightings controlled by the free inverse temperature parameters *β^MB^* and *β^MF^*; a third parameter *β^stick^* captures any value-independent bias to stay or switch. Second-level (alien) choices are determined by a single set of *Q* values (since model-based and model-free evaluation coincide for terminal actions), with inverse temperature *β^stage^*^2^. The various *Q* values are updated according to delta rules with a free learning rate parameter *α.* Finally, the net model-free weighting *β^MF^* is itself derived from the weighted combination of *Q* values learned by two variants of TD learning, TD(0) and TD(1), with weights *β^MF^*^0^ and *β^MF^*^1^. (This is a minor change of variables with respect to the standard model-free TD(λ) algorithm used to hybridize these learning rules in Daw et al., 2011. Here the second temperature parameter replaces the eligibility trace parameter *λ* used in that model, which has the advantage of eliminating its 0,1 boundaries.) Following estimation, we reverse the change of variables by computing the net model-free weighting as 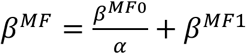, where the *α* accounts for a difference in scaling between the two parameters (see Gillan et al., 2016). When making group comparisons, group estimates of *β^MF^*^0^ are scaled by the estimated *α* of the corresponding group.

The model thus estimates six free parameters per participant: *α,β^MB^, β^MF^*^0^*, β^MF^*^1^*, β^stick^*, and *β^stage^*^2^, and our main hypotheses of interest concern group-wise differences in *β^MB^* and the net *β^MF^*.

### Group-level modeling and estimation

The model was specified hierarchically, so that the participant-specific parameter estimates were assumed to be drawn from a population-level distribution, separately for the patient and control groups. In particular, parameters 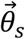 (a six-vector) for each participant *s* were modeled as drawn from a multivariate normal with mean 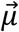 and covariance *Σ.* An additional vector 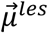 coded any difference in means for the lesion group (i.e., their mean was 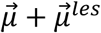, allowing us to test for group differences in each parameter by comparing the corresponding element of 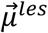 to zero). For the parameter *α* (which is constrained to 0,1), the corresponding element of 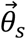 (which has infinite support) was transformed through the CDF of the standard normal.

We jointly estimated the posterior distribution over the individual and group-level parameters using MCMC as described above, which required specifying prior distributions (‘‘hyperpriors’’) on the parameters of the group level distributions. In particular, priors for the elements of 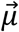 and 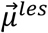 were individually normal (mean=0, SD=2), which is uninformative within the relevant range. The covariance *Σ* was specified (as recommended in the Stan documentation) as the product of a correlation matrix *Ω* (which had an LKJ prior with shape *v* =2; Lewandowski et al., 2009) scaled element wise by the outer product of a scale vector 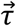 (whose elements were again taken as normal, mean=0, sd=2) with itself. This model also included individual IQ scores and age as covariates.

In order to test the interaction between performance in the two tasks, we then expressed a new model with group-specific parameter estimates (priors were normal distributions with mean = 0 and sd=1) that specified how individual z-scored estimates of place memory (dB predicted the participant-specific parameter estimates. dB was extracted identically as previously. This model also included individual IQ scores as a covariate.

## Supplementary Results from Spatial Task

We compared the behavior of patients and controls in the spatial memory task, which probes the extent to which they can learn locations via an allocentric map vs. via associations with landmarks. A general overview of spatial memory behavior is shown in in Supplementary Figure 1, which plots Distance Error over the 16 presentations of the 4 objects.

**Supplementary Figure 1:**
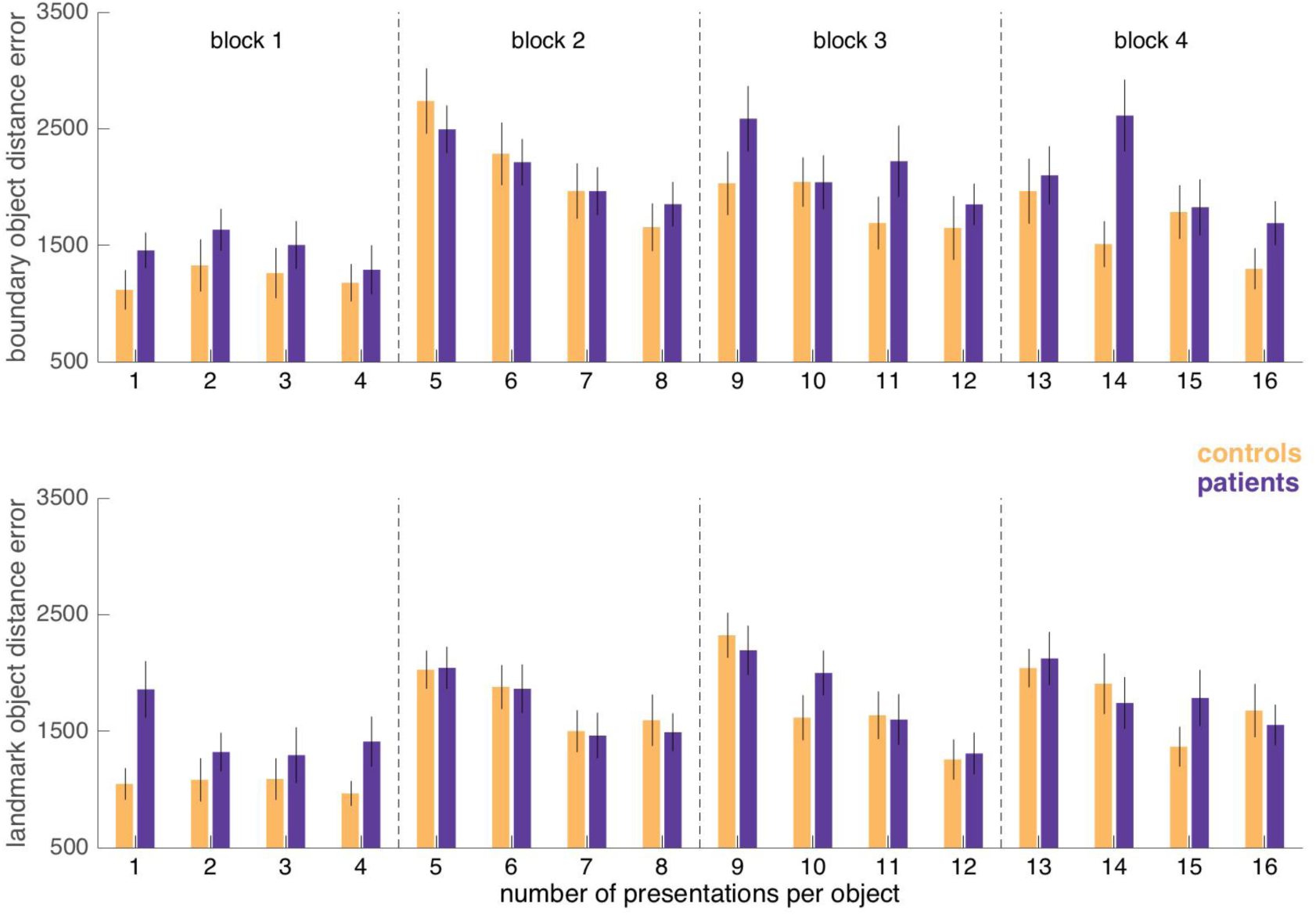
Distance error for boundary objects (Top) and landmark objects (Bottom) over the 16 times that each object is presented. Note that the landmark moves in relation to boundaries before trials object presentation 5, 9 and 13.

As in the earlier fMRI study (Doeller et al., 2008), qualitative differences between performance in the first and remaining three blocks of the task are visible. In the first block (prior to the movement of the landmark disrupting responses), distance error was lower and fairly constant across trials of the block. Distance error was almost significantly greater for patients during this first block (F_1,38_= 4.0279, p=0.05190). The low distance error already from object presentation 1 most likely reflects the fact that the location of each object was shown to participants before they made their first response.

As in Doeller et al. (2008) study, during blocks 2-4, distance errors appeared to spike following the movement of the landmark in relation to the boundaries, and then decrease across the trials within these blocks. The decrease of distance error within blocks 2-4 was significant (F_1, 182_=8.5592, p<0.001), but this reduction did not interact with object type, group, or an interaction of the two. Average distance error was also significantly greater for boundary compared to landmark objects in blocks 2-4 (F_1,38_=8.4522, p=0.006).

Furthermore, while there was no overall or group-wise difference in reduction in distance error across blocks, there was a significantly greater reduction in distance error for boundary objects across blocks (F_1,37_=4.7699 p=0.03529), although this difference did not interact significantly with group (F_1,37_=1.1397, p=0.29255).

## Supplementary Result Tables

**Supplementary Table 1:** Results from from linear mixed regression model, with participant as random factor, implemented using Matlab function fitglme. Dependent variable is Distance Error. Age and IQ are z-scored. Effects seen for Patient group and Control group separately.

| Term | F | DF1 | DF2 | pValue | Estimate | SE |
| --- | --- | --- | --- | --- | --- | --- |
| 'zIQ:dB' | 15.29 | 1 | 39.42 | 0 | -369.1 | 94.42 |
| 'zAge:dB' | 3.455 | 1 | 39.47 | 0.071 | 176.5 | 94.97 |
| 'Controls:dB' | 275.3 | 1 | 39.48 | 0 | 2219 | 133.7 |
| 'Patients:dB' | 406.2 | 1 | 39.51 | 0 | 2695 | 133.7 |
| 'zIQ:dL' | 1.425 | 1 | 85.45 | 0.236 | -86.58 | 72.52 |
| 'zAge:dL' | 1.755 | 1 | 85.55 | 0.189 | -96.65 | 72.95 |
| 'Controls:dL' | 615.8 | 1 | 85.56 | 0 | 2549 | 102.7 |
| 'Patients:dL' | 615.8 | 1 | 85.6 | 0 | 2549 | 102.7 |

**Supplementary Table 2:** Results from from binomial mixed regression model, with participant as random factor, implemented using Matlab function fitglme. Dependent variable is whether participant stay with the first same first level action as on the last trial. Rare transition on last trial coded as 1, common as −1. Rewards on last trials coded as 1, no reward coded as −1. Age and IQ are z-scored. Effects seen for Patient group and Control group separately.

|  | z | pValue | Estimate | SE |
| --- | --- | --- | --- | --- |
| 'Patients' | 7.57 | 0 | 1.635 | 0.2161 |
| 'Controls' | 7.13 | 0 | 1.533 | 0.2152 |
| 'Reward:zAge' | 1.55 | 0.121 | 0.0778 | 0.0502 |
| 'Rare:zAge' | 0.96 | 0.34 | 0.0515 | 0.0539 |
| 'Reward:zIQ' | 0.45 | 0.655 | 0.0234 | 0.0523 |
| 'Rare:zIQ' | 0.16 | 0.872 | -0.0089 | 0.0555 |
| 'Reward:Patients' | 6.05 | 0 | 0.5114 | 0.0845 |
| 'Rare:Patients' | 1.01 | 0.313 | -0.079 | 0.0783 |
| 'Reward:Controls' | 4.41 | 0 | 0.3664 | 0.0831 |
| 'Rare:Controls' | 0.55 | 0.579 | -0.0426 | 0.0767 |
| 'Reward:Rare:zAge' | 1.45 | 0.147 | 0.0828 | 0.0571 |
| 'Reward:Rare:zIQ' | 0.99 | 0.322 | -0.0582 | 0.0587 |
| 'Reward:Rare:Patients' | 1.35 | 0.176 | -0.1158 | 0.0856 |
| 'Reward:Rare:Controls' | 3.32 | 0.001 | -0.2794 | 0.0841 |

**Supplementary Table 3:** Results from from binomial mixed regression model, with participant as random factor, implemented using Matlab function fitglme. Dependent variable is whether participant stay with the first same first level action as on the last trial. Rare transition on last trial coded as 1, common as −1. Rewards on last trials coded as 1, no reward coded as −1. Controlling for z-scored IQ. dB is z-scored participant wise place memory performance. Effects seen for Patient group and Control group separately.

|  | z | pValue | Estimate | SE |
| --- | --- | --- | --- | --- |
| 'Controls' | 6.74 | 0 | 1.505 | 0.2234 |
| 'Reward:zIQ' | 1.04 | 0.296 | 0.065 | 0.0623 |
| 'Rare:zIQ' | 0.62 | 0.533 | 0.042 | 0.0675 |
| ' dB:Patients' | 0.57 | 0.569 | -0.1413 | 0.2482 |
| 'Reward:Patients' | 5.09 | 0 | 0.4601 | 0.0905 |
| 'Rare:Patients' | 0.71 | 0.479 | -0.0579 | 0.0817 |
| ' dB:Controls' | 0.62 | 0.538 | -0.1324 | 0.2151 |
| 'Reward:Controls' | 4.44 | 0 | 0.3964 | 0.0892 |
| 'Rare:Controls' | 0.13 | 0.898 | 0.0104 | 0.0809 |
| 'Reward:Rare:zIQ' | 0.58 | 0.564 | 0.0409 | 0.0708 |
| ' dB:Reward:Patients' | 1.29 | 0.196 | 0.132 | 0.1021 |
| ' dB:Rare:Patients' | 1.05 | 0.293 | -0.098 | 0.0933 |
| 'Reward:Rare:Patients' | 1.46 | 0.145 | -0.1271 | 0.0872 |
| ' dB:Reward:Controls' | 0.19 | 0.848 | 0.0195 | 0.1017 |
| ' dB:Rare:Controls' | 1.56 | 0.119 | 0.1516 | 0.0973 |
| 'Reward:Rare:Controls' | 2.12 | 0.034 | -0.1834 | 0.0866 |
| ' dB:Reward:Rare:Patients' | 0.18 | 0.859 | -0.0176 | 0.0991 |
| ' dB:Reward:Rare:Controls' | 2.66 | 0.008 | 0.2752 | 0.1036 |

**Supplementary Table 4:** Results from from linear mixed regression model, with participant as random factor, implemented using Matlab function fitglme. Dependent variable is Distance Error. Age and IQ are z-scored. Effects seen for Left patient group, Right patient group and Control group separately.

|  | F | DF1 | DF2 | pValue | Estimate | SE |
| --- | --- | --- | --- | --- | --- | --- |
| 'zIQ:dB' | 15.21 | 1 | 39.36 | 0 | -368.4 | 94.44 |
| 'zAge:dB' | 3.232 | 1 | 39.57 | 0.08 | 173.2 | 96.36 |
| 'Controls:dB' | 275.7 | 1 | 39.4 | 0 | 2219 | 133.7 |
| 'Left:dB' | 192.7 | 1 | 41.36 | 0 | 2724 | 196.2 |
| 'Right:dB' | 208.3 | 1 | 38.11 | 0 | 2668 | 184.9 |
| 'zIQ:dL' | 1.35 | 1 | 88.62 | 0.248 | -83.97 | 72.27 |
| 'zAge:dL' | 2.114 | 1 | 89.27 | 0.15 | -107.3 | 73.8 |
| 'Controls:dL' | 621.7 | 1 | 88.67 | 0 | 2550 | 102.3 |
| 'Left:dL' | 304 | 1 | 95.42 | 0 | 2640 | 151.4 |
| 'Right:dL' | 308.5 | 1 | 84.24 | 0 | 2471 | 140.7 |

**Supplementary Table 5:**
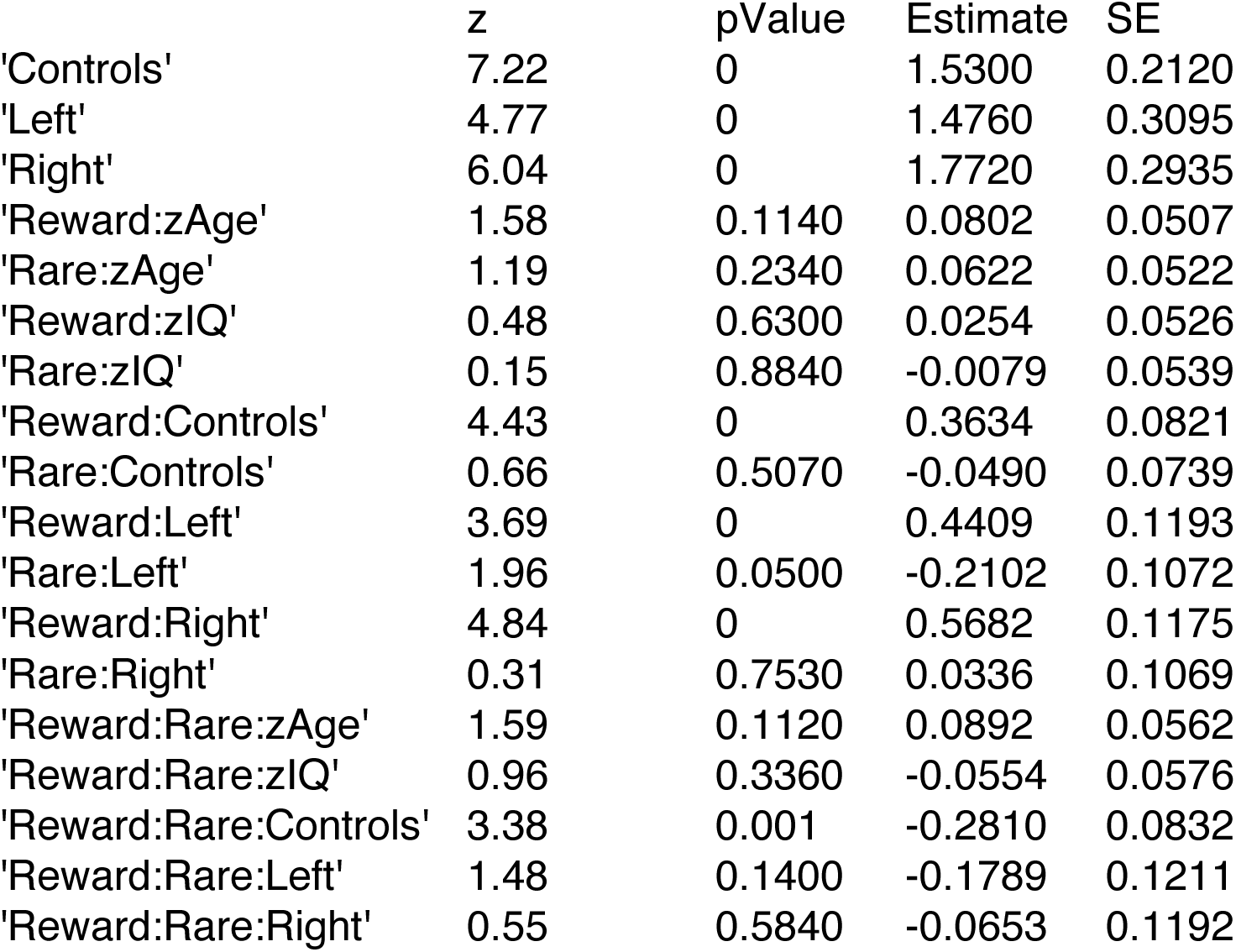
Results from from binomial mixed regression model, with participant as random factor, implemented using Matlab function fitglme. Dependent variable is whether participant stay with the first same first level action as on the last trial. Rare transition on last trial coded as 1, common as −1. Rewards on last trials coded as 1, no reward coded as −1. Age and IQ are z-scored. Effects seen for Left Patient group, Right Patient group and Control group separately.

**Supplementary Table 6:** Results from from binomial mixed regression model, with participant as random factor, implemented using Matlab function fitglme. Dependent variable is whether participant stay with the first same first level action as on the last trial. Rare transition on last trial coded as 1, common as −1. Rewards on last trials coded as 1, no reward coded as −1. IQ and participant-wise dB scores are z-scored. Effects seen for Left Patient group, Right Patient group and Control group separately.

|  | z | pValue | Estimate | SE |
| --- | --- | --- | --- | --- |
| 'Controls' | 6.81 | 0 | 1.4980 | 0.2201 |
| 'Left' | 4.4 | 0 | 1.5000 | 0.3405 |
| 'Right' | 6.02 | 0 | 1.8080 | 0.3002 |
| 'Reward:zIQ' | 0.42 | 0.6720 | 0.0276 | 0.0651 |
| 'Rare:zIQ' | 0.29 | 0.7730 | 0.0201 | 0.0697 |
| 'Reward:Controls' | 4.38 | 0 | 0.3763 | 0.0860 |
| 'Rare:Controls' | 0.04 | 0.9660 | -0.0033 | 0.0778 |
| 'Reward:Left' | 2.41 | 0.0160 | 0.3154 | 0.1309 |
| 'Rare:Left' | 1.69 | 0.0900 | -0.1993 | 0.1177 |
| 'Reward:Right' | 4.67 | 0 | 0.5586 | 0.1196 |
| 'Rare:Right' | 0.41 | 0.6810 | 0.0446 | 0.1086 |
| 'Controls:zdB' | 0.66 | 0.5090 | -0.1400 | 0.2119 |
| 'Left:zdB' | 0.15 | 0.8790 | -0.0523 | 0.3448 |
| 'Right:zdB' | 0.46 | 0.6470 | -0.1631 | 0.3559 |
| 'Reward:Rare:zIQ' | 0.03 | 0.9760 | -0.0022 | 0.0728 |
| 'Reward:Rare:Controls' | 2.42 | 0.0160 | -0.2034 | 0.0841 |
| 'Reward:Rare:Left' | 1.84 | 0.0660 | -0.2340 | 0.1275 |
| 'Reward:Rare:Right' | 0.5 | 0.6160 | -0.0587 | 0.1168 |
| 'Reward:Controls:zdB' | 0.18 | 0.8560 | -0.0182 | 0.0997 |
| 'Rare:Controls:zdB' | 1.32 | 0.1870 | 0.1258 | 0.0954 |
| 'Reward:Left:zdB' | 2.03 | 0.0420 | 0.2899 | 0.1429 |
| 'Rare:Left:zdB' | 0.01 | 0.9890 | 0.0018 | 0.1304 |
| 'Reward:Right:zdB' | 0.02 | 0.9810 | -0.0036 | 0.1468 |
| 'Rare:Right:zdB' | 1.11 | 0.2680 | -0.1500 | 0.1355 |
| 'Reward:Rare:Controls:zdB' | 2.29 | 0.0220 | 0.2332 | 0.1019 |
| 'Reward:Rare:Left:zdB' | 1.01 | 0.3150 | 0.1402 | 0.1394 |
| 'Reward:Rare:Right:zdB' | 1.17 | 0.2430 | -0.1698 | 0.1453 |

**Supplementary Table 7:** Results from from linear mixed regression model, with participant as random factor, implemented using Matlab function fitglme. Dependent variable is Boundary Distance Error. Age, IQ Total Lesion size and Hippocampal Lesion size are z-scored. Effects seen for Left Patient group and Right Patient group separately.

|  | F | DF1 | DF2 | pValue | Estimate | SE |
| --- | --- | --- | --- | --- | --- | --- |
| 'zIQ' | 1.0836 | 1 | 11.706 | 0.319 | 0.1299 | 0.1247 |
| 'zAge' | 0.7621 | 1 | 11.993 | 0.4 | -0.1088 | 0.1247 |
| 'Left' | 0.5490 | 1 | 13.236 | 0.472 | 0.1285 | 0.1736 |
| 'Right' | 0.1521 | 1 | 11.944 | 0.703 | -0.0624 | 0.1599 |
| 'TotalLesion' | 2.6406 | 1 | 12.481 | 0.129 | 0.2826 | 0.1710 |
| 'Left:HippLesion' | 0.1204 | 1 | 12.014 | 0.734 | 0.0686 | 0.1975 |
| 'Right:HippLesion' | 2.6082 | 1 | 12.158 | 0.132 | -0.2947 | 0.1825 |

**Supplementary Table 8:** Results from from binomial mixed regression model, with participant as random factor, implemented using Matlab function fitglme. Dependent variable is whether participant stay with the first same first level action as on the last trial. Rare transition on last trial coded as 1, common as −1. Rewards on last trials coded as 1, no reward coded as −1. Age, IQ Total Lesion size and Hippocampal Lesion size are z-scored. Effects seen for Left Patient group and Right Patient group separately.

|  | z | pValue | Estimate | SE |
| --- | --- | --- | --- | --- |
| 'Left' | 5.63 | 0 | 1.605 | 0.2852 |
| 'Right' | 7.31 | 0 | 1.964 | 0.2686 |
| 'Reward:zAge' | 1.64 | 0.102 | 0.1481 | 0.0906 |
| 'Rare:zAge' | 0.42 | 0.673 | -0.0369 | 0.0874 |
| 'Reward:zIQ' | 0.25 | 0.8 | -0.024 | 0.0947 |
| 'Rare:zIQ' | 1.71 | 0.086 | 0.1529 | 0.0892 |
| 'Reward:Left' | 2.96 | 0.003 | 0.3868 | 0.1308 |
| 'Rare:Left' | 1.92 | 0.055 | -0.2367 | 0.1234 |
| 'Reward:Right' | 5.43 | 0 | 0.6958 | 0.1281 |
| 'Rare:Right' | 0.58 | 0.564 | 0.0708 | 0.1226 |
| 'Left:HippLesion' | 1.69 | 0.091 | 0.4991 | 0.2956 |
| 'Right:HippLesion' | 2.2 | 0.028 | -0.5592 | 0.2542 |
| 'Reward:TotalLesion' | 0.14 | 0.885 | 0.0177 | 0.1225 |
| 'Rare:TotalLesion' | 2.29 | 0.022 | -0.2682 | 0.1173 |
| 'Reward:Rare:zAge' | 0.7 | 0.487 | 0.0671 | 0.0965 |
| 'Reward:Rare:zIQ' | 0.94 | 0.347 | 0.094 | 0.0999 |
| 'Reward:Rare:Left' | 2.51 | 0.012 | -0.3236 | 0.1287 |
| 'Reward:Rare:Right' | 0.43 | 0.664 | -0.0552 | 0.1269 |
| 'Reward:Left:HippLesion' | 0.65 | 0.519 | -0.088 | 0.1364 |
| 'Rare:Left:HippLesion' | 1.63 | 0.102 | 0.2078 | 0.1271 |
| 'Reward:Right:HippLesion' | 1.83 | 0.067 | -0.2489 | 0.1359 |
| 'Rare:Right:HippLesion' | 1.75 | 0.08 | 0.2267 | 0.1295 |
| 'Reward:Rare:TotalLesion' | 2.32 | 0.02 | -0.3008 | 0.1297 |
| 'Reward:Rare:Left:HippLesion' | 0.11 | 0.912 | -0.0149 | 0.1354 |
| 'Reward:Rare:Right:HippLesion' | 2.82 | 0.005 | 0.385 | 0.1367 |

**Supplementary Table 9:** Results from from binomial mixed regression model, with participant as random factor, implemented using Matlab function fitglme. Dependent variable is whether participant stay with the first same first level action as on the last trial. Rare transition on last trial coded as 1, common as −1. Rewards on last trials coded as 1, no reward coded as −1. IQ, Age, and Post-Rare Slowing are z-scored.

|  | z | pValue | Estimate | SE |
| --- | --- | --- | --- | --- |
| '(Intercept)' | 11.45 | 0 | 1.5900 | 0.1389 |
| 'Reward' | 7.54 | 0 | 0.4438 | 0.0588 |
| 'Rare' | 1.28 | 0.2000 | -0.0661 | 0.0516 |
| 'zAge' | 1.63 | 0.1040 | 0.2442 | 0.1502 |
| 'zIQ' | 0.06 | 0.9540 | 0.0089 | 0.1534 |
| 'zRareSlow' | 2.34 | 0.0190 | 0.3781 | 0.1614 |
| 'Reward:Rare' | 3.78 | 0 | -0.1986 | 0.0525 |
| 'Reward:zAge' | 2.08 | 0.0380 | 0.1299 | 0.0625 |
| 'Rare:zAge' | 0.18 | 0.8550 | 0.0100 | 0.0543 |
| 'Reward:zIQ' | 0.23 | 0.8150 | 0.0152 | 0.0648 |
| 'Rare:zIQ' | 0.68 | 0.4940 | 0.0387 | 0.0567 |
| 'Reward:zRareSlow' | 1.46 | 0.1440 | 0.1020 | 0.0699 |
| 'Rare:zRareSlow' | 2.13 | 0.0330 | -0.1320 | 0.0618 |
| 'Reward:Rare:zAge' | 0.34 | 0.7330 | -0.0190 | 0.0555 |
| 'Reward:Rare:zIQ' | 0.26 | 0.7920 | 0.0152 | 0.0578 |
| 'Reward:Rare:zRareSlow' | 3.53 | 0 | -0.2226 | 0.0630 |

**Supplementary Table 10:** Results from from binomial mixed regression model, with participant as random factor, implemented using Matlab function fitglme. Dependent variable is whether participant stay with the first same first level action as on the last trial. Rare transition on last trial coded as 1, common as −1. Rewards on last trials coded as 1, no reward coded as −1. IQ, Age, and Post-Rare Slowing are z-scored. Effects seen for Left Patient group, Right Patient group and Control group separately.

|  | F | pValue | Estimate | SE |
| --- | --- | --- | --- | --- |
| 'Controls' | 7.67 | 0 | 1.535 | 0.2003 |
| 'Left' | 4.39 | 0 | 1.524 | 0.3469 |
| 'Right' | 6.04 | 0 | 1.735 | 0.2874 |
| 'Reward:zIQ' | 0.35 | 0.729 | 0.02 | 0.0577 |
| 'Rare:zIQ' | 0.96 | 0.336 | 0.0487 | 0.0507 |
| 'Reward:Controls' | 4.5 | 0 | 0.3795 | 0.0843 |
| 'Rare:Controls' | 0.7 | 0.485 | -0.045 | 0.0645 |
| 'Reward:Left' | 3.51 | 0 | 0.5198 | 0.148 |
| 'Rare:Left' | 3.13 | 0.002 | -0.3557 | 0.1135 |
| 'Reward:Right' | 4.5 | 0 | 0.5528 | 0.1227 |
| 'Rare:Right' | 0.38 | 0.701 | 0.0365 | 0.0949 |
| 'Controls:zRareSlow' | 2.02 | 0.043 | 0.3344 | 0.1651 |
| 'Left:zRareSlow' | 0.28 | 0.781 | 0.1573 | 0.5666 |
| 'Right:zRareSlow' | 0.44 | 0.661 | 0.151 | 0.3439 |
| 'Reward:Rare:zIQ' | 0.01 | 0.993 | 0 | 0.0541 |
| 'Reward:Rare:Controls' | 3.95 | 0 | -0.2723 | 0.069 |
| 'Reward:Rare:Left' | 2.65 | 0.008 | -0.3222 | 0.1214 |
| 'Reward:Rare:Right' | 0.43 | 0.667 | -0.0436 | 0.1013 |
| 'Reward:Controls:zRareSlow' | 0.87 | 0.383 | 0.0662 | 0.0759 |
| 'Rare:Controls:zRareSlow' | 2.55 | 0.011 | -0.1543 | 0.0604 |
| 'Reward:Left:zRareSlow' | 0.85 | 0.396 | 0.1996 | 0.2353 |
| 'Rare:Left:zRareSlow' | 2.45 | 0.014 | -0.4376 | 0.1785 |
| 'Reward:Right:zRareSlow' | 0.53 | 0.599 | -0.0795 | 0.151 |
| 'Rare:Right:zRareSlow' | 0.38 | 0.705 | -0.0458 | 0.121 |
| 'Reward:Rare:Controls:zRareSlow' | 3.14 | 0.002 | -0.2023 | 0.0645 |
| 'Reward:Rare:Left:zRareSlow' | 2.42 | 0.016 | -0.4627 | 0.1912 |
| 'Reward:Rare:Right:zRareSlow' | 1.7 | 0.09 | -0.2169 | 0.1277 |

**Supplementary Table 11:** Results from from linear mixed regression model, with participant as random factor, implemented using Matlab function fitglme. Dependent variable is Post-Rare Slowing. Age, IQ, Total Lesion size and Hippocampal Lesion size are z-scored. Effects seen for Left Patient group and Right Patient group separately.

|  | F | DF1 | DF2 | pValue | Estimate | SE |
| --- | --- | --- | --- | --- | --- | --- |
| 'zIQ' | 1.4 | 1 | 19.12 | 0.251 | -0.0478 | 0.0404 |
| 'Left' | 6.665 | 1 | 19.16 | 0.018 | 0.1353 | 0.0524 |
| 'Right' | 0.0525 | 1 | 19.01 | 0.821 | 0.0112 | 0.049 |
| 'TotalLesion' | 7.955 | 1 | 19.14 | 0.011 | 0.1498 | 0.0531 |
| 'Left:HippLesion' | 2.723 | 1 | 18.99 | 0.115 | -0.1042 | 0.0632 |
| 'Right:HippLesion' | 1.825 | 1 | 19 | 0.193 | -0.0753 | 0.0557 |
| 'zAge:Rare' | 12.87 | 1 | 19.66 | 0.002 | -0.0309 | 0.0086 |
| 'zIQ:Rare' | 0.7316 | 1 | 20.8 | 0.402 | 0.0075 | 0.0088 |
| 'Left:Rare' | 61.37 | 1 | 21.97 | 0 | 0.0912 | 0.0116 |
| 'Right:Rare' | 58.89 | 1 | 18.5 | 0 | 0.0799 | 0.0104 |
| 'TotalLesion:Rare' | 9.336 | 1 | 21.71 | 0.006 | 0.0358 | 0.0117 |
| 'Left:HippLesion:Rare' | 0.0902 | 1 | 17.89 | 0.767 | 0.004 | 0.0133 |
| 'Right:HippLesion:Rare' | 9.993 | 1 | 18.69 | 0.005 | -0.0375 | 0.0119 |

**Supplementary Table 12:** Estimates (mean, 2.5% confidence interval and 97.5% confidence interval) from the full RL model which includes age and IQ as covariates but not individual place-memory performance estimates. Cont indicated free-parameter estimates of the control group. Pat_diff indicates how patient group free-parameter estimates differ from control group.

|  | mean | 2.5% | 97.5% |
| --- | --- | --- | --- |
| $\beta_{\text{cont}}^{\text{MB}}$ | 0.33 | 0.1 | 0.55 |
| $\beta_{\text{cont}}^{\text{MF0}}$ | -0.06 | -0.14 | 0.01 |
| $\beta_{\text{cont}}^{\text{MF1}}$ | 0.30 | 0.15 | 0.46 |
| $\beta_{\text{cont}}^{\text{stick}}$ | 1.16 | 0.78 | 1.54 |
| $\beta_{\text{cont}}^{2\text{stage}}$ | 1.17 | 0.85 | 1.5 |
| $\alpha_{\text{cont}}$ | 0.26 | 0.18 | 0.38 |
| $\beta_{\text{pat\_diff}}^{\text{MB}}$ | -0.2 | -0.52 | 0.13 |
| $\beta_{\text{pat\_diff}}^{\text{MF0}}$ | 0.05 | -0.05 | 0.15 |
| $\beta_{\text{pat\_diff}}^{\text{MF1}}$ | 0.2 | -0.02 | 0.42 |
| $\beta_{\text{pat\_diff}}^{\text{stick}}$ | 0.25 | -0.29 | 0.79 |
| $\beta_{\text{pat\_diff}}^{2\text{stage}}$ | -0.04 | -0.49 | 0.41 |
| $\alpha_{\text{pat\_diff}}$ | 0.07 | -0.06 | 0.22 |

**Supplementary Table 13:** Estimates (mean, 2.5% confidence interval and 97.5% confidence interval) from the full RL model which includes IQ and individual place memory performance estimates as covariates. Cont indicates effect of place-memory (dB) on free-parameter estimates of the control group. Pat_diff indicates how patient group effect of place-memory on free parameter estimates differ from control group.

|  | mean | 2.5% | 97.5% |
| --- | --- | --- | --- |
| $\beta^{\text{MB-dB}}_{\text{cont}}$ | -0.27 | -0.52 | -0.02 |
| $\beta^{\text{MF0-dB}}_{\text{cont}}$ | 0.00 | -0.08 | 0.07 |
| $\beta^{\text{MF1-dB}}_{\text{cont}}$ | 0.12 | -0.07 | 0.31 |
| $\beta^{\text{stick-dB}}_{\text{cont}}$ | -0.13 | -0.59 | 0.32 |
| $\beta^{\text{2stage-dB}}_{\text{cont}}$ | -0.38 | -0.73 | -0.03 |
| $\alpha\text{-dB}_{\text{cont}}$ | 0.24 | 0.15 | 0.34 |
| $\beta^{\text{MB-dB}}_{\text{pat\_diff}}$ | 0.29 | -0.05 | 0.63 |
| $\beta^{\text{MF0-dB}}_{\text{pat\_diff}}$ | -0.03 | -0.15 | 0.09 |
| $\beta^{\text{MF1-dB}}_{\text{pat\_diff}}$ | -0.05 | -0.31 | 0.21 |
| $\beta^{\text{stick-dB}}_{\text{pat\_diff}}$ | 0.06 | -0.54 | 0.66 |
| $\beta^{\text{2stage-dB}}_{\text{pat\_diff}}$ | 0.51 | 0.03 | 0.97 |
| $\alpha\text{-dB}_{\text{pat\_diff}}$ | 0.06 | -0.06 | 0.22 |

